# An anti-SARS-CoV-2 non-neutralizing antibody with Fc-effector function defines a new NTD epitope and delays neuroinvasion and death in K18-hACE2 mice

**DOI:** 10.1101/2021.09.08.459408

**Authors:** Guillaume Beaudoin-Bussières, Yaozong Chen, Irfan Ullah, Jérémie Prévost, William D. Tolbert, Kelly Symmes, Shilei Ding, Mehdi Benlarbi, Shang Yu Gong, Alexandra Tauzin, Romain Gasser, Debashree Chatterjee, Dani Vézina, Guillaume Goyette, Jonathan Richard, Fei Zhou, Leonidas Stamatatos, Andrew T. McGuire, Hughes Charest, Michel Roger, Edwin Pozharski, Priti Kumar, Walther Mothes, Pradeep D. Uchil, Marzena Pazgier, Andrés Finzi

## Abstract

Emerging evidence in animal models indicate that both neutralizing activity and Fc-mediated effector functions of neutralizing antibodies contribute to protection against SARS-CoV-2. It is unclear if antibody effector functions alone could protect against SARS-CoV-2. Here we isolated CV3-13, a non-neutralizing antibody from a convalescent individual with potent Fc-mediated effector functions that targeted the N- terminal domain (NTD) of SARS-CoV-2 Spike. The cryo-EM structure of CV3-13 in complex with SAR-CoV-2 spike revealed that the antibody bound from a distinct angle of approach to a novel NTD epitope that partially overlapped with a frequently mutated NTD supersite in SARS-CoV-2 variants. While CV3-13 did not alter the replication dynamics of SARS-CoV-2 in a K18-hACE2 transgenic mouse model, an Fc-enhanced CV3-13 significantly delayed neuroinvasion and death in prophylactic settings. Thus, we demonstrate that efficient Fc-mediated effector functions can contribute to the *in vivo* efficacy of anti-SARS-CoV-2 monoclonal antibodies in the absence of neutralization.

## Introduction

In December of 2019, a new coronavirus was detected in Wuhan, China which has subsequently been named severe acute respiratory syndrome coronavirus 2 (SARS-CoV-2) (Huang et al., 2020; Wu et al., 2020b; Zhu et al., 2020). This novel virus is an enveloped, positive-strand RNA coronavirus that is a member of the β-coronavirus genus (Huang et al., 2020; Wu et al., 2020b; Zhu et al., 2020). This genus also contains SARS-CoV-1 and MERS-CoV which caused pandemics in 2002 and 2012, respectively (Ksiazek et al., 2003; Zaki et al., 2012). SARS-CoV-2, however, has caused a much larger pandemic with millions of deaths worldwide (Worldometer, 2021). The efficacy with which SARS-CoV-2 transmits between humans has led to COVID-19 being one of the largest and fastest growing pandemics in the past century. The COVID-19 pandemic has caused a massive loss of human life, extreme economic consequences and disrupted billions of human lives worldwide. For these reasons, billions of dollars have been invested in the development of vaccines against SARS-CoV-2 which has resulted in their development in record time. Several vaccine platforms have been approved in different jurisdictions worldwide to counter the COVID-19 pandemic (Oliver et al., 2021a; Oliver et al., 2021b; Polack et al., 2020; Voysey et al., 2021) with vaccine development primarily focused on generating immune responses against the SARS-CoV-2 Spike glycoprotein. The Spike glycoprotein mediates viral entry and is well exposed at the surface of virions (Duan et al., 2020; Shang et al., 2020) and infected cells (Buchrieser et al., 2020). The Spike glycoprotein is a trimer of heterodimers, composed of 2 subunits, S1 and S2, generated by furin cleavage of a single S polypeptide. The S1 subunit permits attachment via its receptor binding domain (RBD) to the cellular receptor angiotensin converting enzyme 2 (ACE2) (Hoffmann et al., 2020; Lan et al., 2020; Walls et al., 2020; Wrapp et al., 2020b). It also contains an N-terminal domain (NTD) that may aid attachment and conformational transition of Spike, as previously observed for other coronaviruses. (Amraie et al., 2020; Lempp et al., 2021; Soh et al., 2020). Once bound to ACE2, a series of conformational changes allows the S2 subunit to mediate fusion between the cellular and viral membranes. Virus neutralization has been shown to be important for controlling SARS-CoV-2 infection in vaccinated human cohorts (AstraZeneca and Iqvia Pty, 2021; Bio and Pfizer, 2021; Janssen and Prevention, 2021; ModernaTx et al., 2022). As a result, considerable effort has been made to study antibody-mediated neutralization and its effect in mitigating SARS-CoV-2 infection. Many neutralizing antibodies target the RBD, but some targeting the NTD and the S2 subunits have also been described (Cao et al., 2020; Chen et al., 2020; Chi et al., 2020; Jennewein et al., 2021; Ju et al., 2020; Li et al., 2021b; Liu et al., 2020; Rappazzo et al., 2021; Seydoux et al., 2020; Suryadevara et al., 2021; Ullah et al., 2021; Voss et al., 2021; Wang et al., 2020; Wrapp et al., 2020a; Wu et al., 2020c; Yuan et al., 2020). However, some studies have shown that around 25% to 45% of people who resolve the infection have plasma with low or undetectable levels of SARS-CoV-2 neutralizing activity (Beaudoin-Bussieres et al., 2020; Luchsinger et al., 2020; Muecksch et al., 2021; Payne et al., 2020; Prevost et al., 2020; Robbiani et al., 2020; Wu et al., 2020a). These data suggest that immune functions other than neutralization could play a role in SARS-CoV- 2 control. Accordingly, results from a phase III clinical trials show a vaccine efficacy of >90% starting 14 days after the injection of a single dose of a BNT162b2 mRNA vaccine, a time at which neutralizing activity from the vaccine is weak (Baden et al., 2021; Polack et al., 2020; Skowronski and De Serres, 2021; Tauzin et al., 2021). Similarly, recent studies have shown that despite a significant loss in neutralizing activity against the B.1.1.7 (Alpha) and B.1.351 (Beta) variants, the AstraZeneca and Pfizer/BioNTech vaccines remain efficacious against these variants (Emary et al., 2021) (Pfizer/BioNTech, 2021).

The antigen binding domain (Fab) of antibodies is critical for neutralization but the crystallizable fragment (Fc) of the antibody can contribute significantly to their *in vivo* efficacy (Bournazos et al.,2019; Bournazos et al., 2014; DiLillo et al., 2014). Fc engagement of Fc gamma receptors (FcγRs) elicits complement-dependent cytotoxicity (CDC), antibody-dependent cellular cytotoxicity (ADCC) and antibody-dependent cellular phagocytosis (ADCP) depending on the effector cell to which they bind. We previously examined the protection mediated by neutralizing antibodies (NAbs) targeting the SARS-CoV-2 Spike in a K18-hACE2 mouse model and the effect of wild-type or Fc-mutated versions of these NAbs in a prophylactic or therapeutic setting (Ullah et al., 2021). The Fc mutations (L234A-L235A, also referred to as LALA), significantly diminished the affinity of antibodies to FcγRIIIa and also impacted Fc-mediated effector functions. Interestingly, in this study we showed LALA mutations significantly diminished the capacity of NAbs to protect mice from a lethal SARS-CoV-2 challenge (Ullah et al., 2021). Similarly, two other studies examining humoral responses in acutely infected individuals found that Fc-mediated effector functions were associated with survival (Brunet-Ratnasingham et al., 2021; Zohar et al., 2020). Therefore, while antibody-mediated neutralization was required for protection, it was not sufficient for viral control.

Serological analysis of the plasma or serum from SARS-CoV-2-infected individuals from multiple sources (Harvey et al., 2021; McCallum et al., 2021; Piccoli et al., 2020) revealed that ∼65-80% of the neutralizing response was from RBD-specific antibodies, with a smaller portion targeting the NTD (∼6-20%) or the S2 subunit (4-20%). Although the glycan-shielded NTD has been proposed to have limited immunogenicity as compared to RBD, NTD-directed antibodies are regarded as a major driving force in imposing selection pressure against the virus eliciting many NTD escape mutations and deletions in emerging SARS-CoV-2 variants (McCallum et al., 2021). Notably, NTD-directed monoclonal NAbs whose structures have been determined recognized a similar glycan-free epitope (Cerutti et al., 2021; Chi et al., 2020; Liu et al., 2020; McCallum et al., 2021; Sun et al., 2021; Voss et al., 2021), named the NTD-supersite (residue 14-20, 140-158 and 245-264) (Harvey et al., 2021; McCallum et al., 2021). There are a greater number of mutations within the NTD supersite than in the NTD scaffold that highlights its importance to the virus (McCarthy et al., 2021). Given that the NTD-supersite-directed NAbs do not interfere with the ACE2-RBD interaction or the shedding of the S1 subunit, their mechanism of neutralization is yet to be determined. Possible mechanisms could include interactions with co-receptors, proteolytic activation by TMPRSS2, or interference with membrane-fusion.

Here we sought to test if Fc-mediated effector functions of antibodies alone could mediate virological control in a lethal K18-hACE2 transgenic mouse model of SARS-CoV-2 using CV3-13, a non-neutralizing antibody with potent Fc-effector functions. Our cryo-EM structure revealed that CV3-13 binds to a novel NTD epitope that partially overlaps with the NTD supersite with a distinct angle of approach, adding to our understanding of how fine epitope specificity and the mode of antibody binding can contribute to antibody function. Several recurrent NTD mutations outside of the NTD supersite associated with immune resistance are found within the CV3-13 epitope, e.g. the N2 loop and the newly identified N2-3 hairpin, suggesting that the epitope coincides with those of other NTD-binding antibodies. While administration of CV3-13 under both prophylactic and therapeutic regimens did not change the replication dynamics of SARS- CoV-2 in a K18-hACE2 transgenic mouse model, an Fc-enhanced version of CV3-13 significantly delayed neuroinvasion and death from a lethal challenge of the virus in a prophylactic setting. Thus, using a NTD-binding non-neutralizing antibody, we demonstrate the contribution of Fc-effector functions in mitigating viral-induced pathogenesis.

## Results

### CV3-13 binds the NTD of the Spike glycoprotein

To test if non-neutralizing antibodies alone can protect from SARS-CoV-2 infection, we characterized a non-neutralizing antibody isolated from the peripheral blood mononuclear cells (PBMCs) of a convalescent individual (CV3) 6 weeks after the onset of symptoms. Using fluorescent SARS-CoV-2 Spike 2P as a probe, we sorted four hundred thirty-two antigen-specific B cells from this donor PBMCs. We successfully generated twenty-seven monoclonal antibodies and tested their ability to neutralize pseudoviral particles carrying Spike protein. CV3-13 bound to Spike but did not neutralize SARS-CoV-2 pseudovirus (Jennewein et al., 2021). To determine the epitope recognized by CV3-13, we analyzed its ability to bind different Spike variants on the surface of transfected cells. CV3-13 efficiently bound the WT (Wuhan-Hu-1 reference strain) and the D614G variant but did not recognize the Spike from the B.1.1.7 (alpha) variant (Figure 1A, B). We took advantage of this differential binding capacity to determine the epitope of CV3-13 by sequentially introducing B.1.1.7 variant mutations into the WT Spike. CV3-13 recognized all but the Δ144 mutant that had a single amino acid deletion located in the S1 NTD (Figure 1B). In agreement with our cell surface binding analyses, surface plasmon resonance (SPR) showed that CV3-13 binds to the SARS-CoV-2 S1 subunit (Figure 1C). Monovalent CV3-13 Fab bound to the stabilized Spike trimer ectodomain (Spike-6P) with nanomolar affinity (K_D_ = ∼55 nM) (Figure 1D).

**Figure 1.**
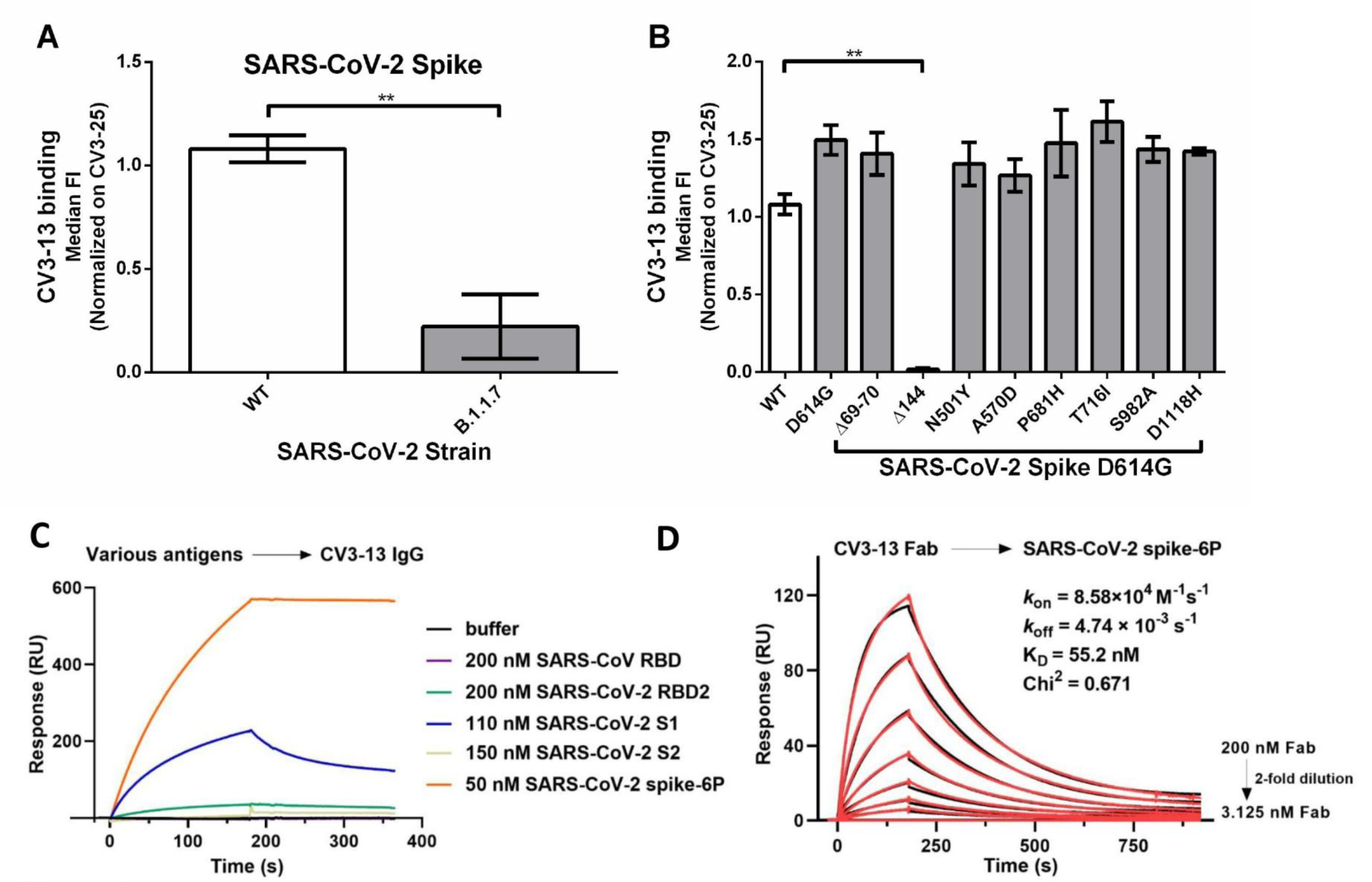
Recognition of SARS-CoV-2 Spikes by CV3-13. (A) Staining of CV3-13 (5 µg/mL) on the Spike of the WT (Wuhan-Hu-1) or the B.1.1.7 (alpha variant) strain of SARS-CoV-2 expressed at the surface of 293T transfected cells. (B) Staining of CV3-13 (5 µg/mL) on the different individual mutations of the Spike of the B.1.1.7 strain of SARS-CoV-2 (D614G, Δ69-70, Δ144, N501Y, A570D, P681H, T716I, S982A and D1118H). CV3-13 binding was further normalized to the binding obtained with the conformational-independent CV3-25 mAb (5 µg/mL). Statistical significance was evaluated using a non-parametric Mann-Whitney U test (**, p < 0.01). Data are the average of the median of each experiment done at least 2 times. Mean values ± Standard error of the mean (SEM). (C) SPR-based epitope mapping reveals that CV3-13 explicitly binds to SARS-CoV-2 spike S1 subunit. Different viral antigens from SARS-CoV or SARS-CoV-2 were injected to the immobilized CV3-13 IgG (∼5800 RU) at the indicated concentrations. (D) Kinetics measurement of CV3-13 Fab binding to the immobilized SARS-CoV-2 HexaPro spike (∼800 RU) with concentrations ranging from 3.125 to 200 nM (2-fold serial dilution). The experimental data (red) were fitted to a 1:1 Langmuir model (black) in BIA evaluation software.

### CV3-13 is a non-neutralizing antibody with potent Fc-mediated effector functions

To confirm that CV3-13 was a non-neutralizing antibody (Jennewein et al., 2021), we tested its capacity to neutralize pseudoviruses carrying the SARS-CoV-2 Spike. For comparison purposes, we used CV3-1, a potent RBD-targeting NAb, as a positive control (Jennewein et al., 2021). CV3-1 was recently shown to protect K18-hACE2 mice from a lethal SARS-CoV-2 challenge in a Fc-effector function-dependent manner (Ullah et al., 2021). Our analyses showed that CV3-13 was unable to neutralize pseudoviral particles (Figure 2A) bearing SARS-CoV-2 Spike or the live virus (Figure 2B). Introduction of LALA or GASDALIE mutations to the Fc portion of the antibody did not modify its neutralization profile (Figure 2A) or its ability to recognize the Spike whether it be at the surface of infected cells, transduced cells or the recombinant SARS-CoV-2 S-6P (Figure 2C, D, S1). The LALA (L234A/L235A) mutations impair the interaction between the IgG Fc portion and FcγRs (Saunders, 2019), while GASDALIE (G236A/S239D/A330L/I332E) mutations strengthen these interactions (Bournazos et al., 2014; DiLillo and Ravetch, 2015; Lazar et al., 2006; Richards et al., 2008; Smith et al., 2012). Having established that CV3-13 does not mediate neutralization (Figure 2A, B), we then evaluated whether it could mediate Fc-effector functions. To this end we used an antibody-dependent cellular cytotoxicity (ADCC) assay using a human T-lymphoid cell line resistant to NK cell-mediated cell lysis (CEM.NKr) and stably expressing the full-length Spike on their surface as target cells. PBMCs from healthy individuals were used as effector cells, as previously reported (Anand et al., 2021). In agreement with a previous study, CV3-1 mediated robust ADCC (Ullah et al., 2021) (Figure 2E). CV3-13 mediated ADCC to a similar level as CV3-1 (Figure 2E). Introduction of the GASDALIE mutations enhanced CV3-13-mediated ADCC to levels surpassing those achieved with CV3-1 or CV3-13 WT at higher concentrations (Figure 2E). As expected, introduction of the LALA mutations significantly decreased CV3-13-mediated ADCC (Figure 2E). CV3- 13 GASDALIE mediated similar ADCP as its WT counterpart, while CV3-13 LALA mediated reduced ADCP (at all tested concentrations) than both CV3-13 WT and CV3-13 GASDALIE (Figure 2F). Altogether, these results confirm that CV3-13 is a non-neutralizing antibody able to mediate Fc-mediated effector functions.

**Figure 2.**
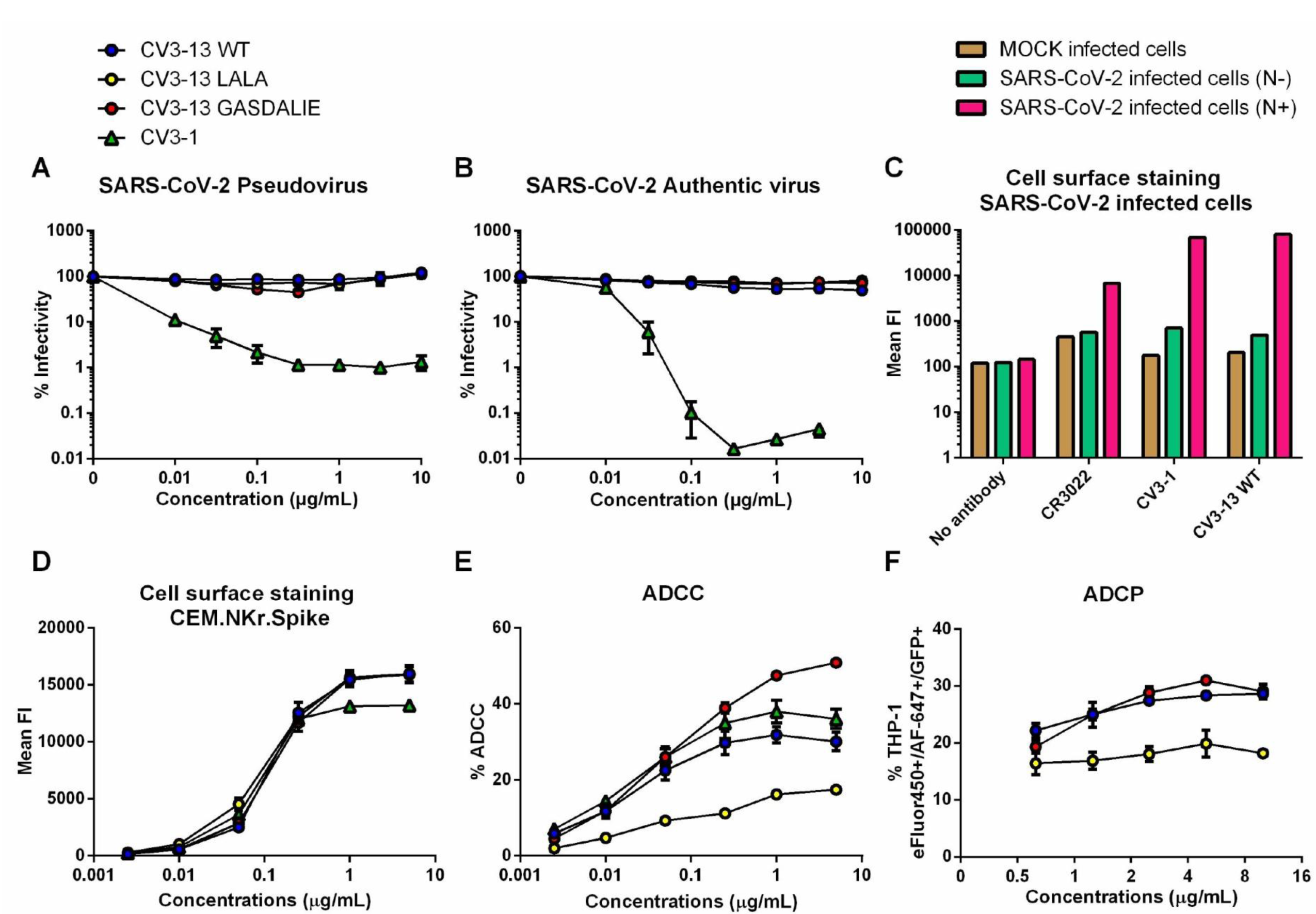
CV3-13 is a non-neutralizing antibody that has potent Fc-mediated effector functions. (A) Neutralizing activity of titrated concentrations of CV3-13 WT, CV3-13 LALA, CV3- 13 GASDALIE and CV3-1 on SARS-CoV-2 Spike D614G bearing pseudoviruses using 293T-ACE2 target cells. (B) Neutralizing activity of CV3-13 WT, CV3-13 LALA, CV3-13 GASDALIE and CV3-1 on SARS-CoV-2 D614G authentic virus using Vero E6 target cells. (C) Binding of CR3022, CV3-1, and CV3-13 WT on the surface of Vero E6 cells infected with authentic SARS-CoV-2 virus 48 hours post-infection. Intracellular Nucleocapsid (N) staining was done to separate infected from the uninfected cells. (D) Binding of CV3-13 WT, CV3-13 LALA, CV3-13 GASDALIE and CV3-1 on CEM.NKr.Spike. (E) % ADCC in the presence of titrated amounts of CV3-13 WT, CV3- 13 LALA, CV3-13 GASDALIE and CV3-1 using a 1:1 ratio of parental CEM.NKr cells and CEM.NKr-Spike cells as target cells while PBMCs from uninfected donors were used as effector cells. (F) % ADCP in the presence of titrated amounts of CV3-13 WT, CV3-13 LALA, CV3-13 GASDALIE and CV3-1 using CEM.NKr-Spike cells as target cells and THP-1 cells as phagocytic cells. Data are the average of at least 2 experiments for (A), (B), (D)-(F). Data is from a single experiment for (C). Mean values ± SEM.

### Structural analyses of the CV3-13 Fab and SARS-CoV-2 spike complex defines a novel NTD epitope

To define the CV3-13 epitope and gain a more comprehensive understanding of how CV3-13 triggers potent Fc-mediated cytotoxicity without directly neutralizing the SARS-CoV-2 virus, we determined the cryo-EM structure of SARS-CoV-2 HexaPro Spike in complex with CV3-13 Fab at a resolution of 4.45 Å (FSC cut-off 0.143) using C1- symmetry (Figure 3 and S4). The Spike trimer presented itself in the one-RBD-up conformation with each of the N-terminal domain (NTD) bound to one CV3-13 Fab to the furthest lateral side of the spike relative to the spike-trimer axis. Despite the asymmetric RBD conformation in the three spike protomers, the three CV3-13-NTD interfaces were identical. Global refinement imposed with C3 symmetry improved the overall resolution to 4.19 Å and permitted a detailed analysis of the CV3-13 footprint. The C3-symmetry map was therefore used in all further structural analysis (Figure 3 and S4).

**Figure 3.**
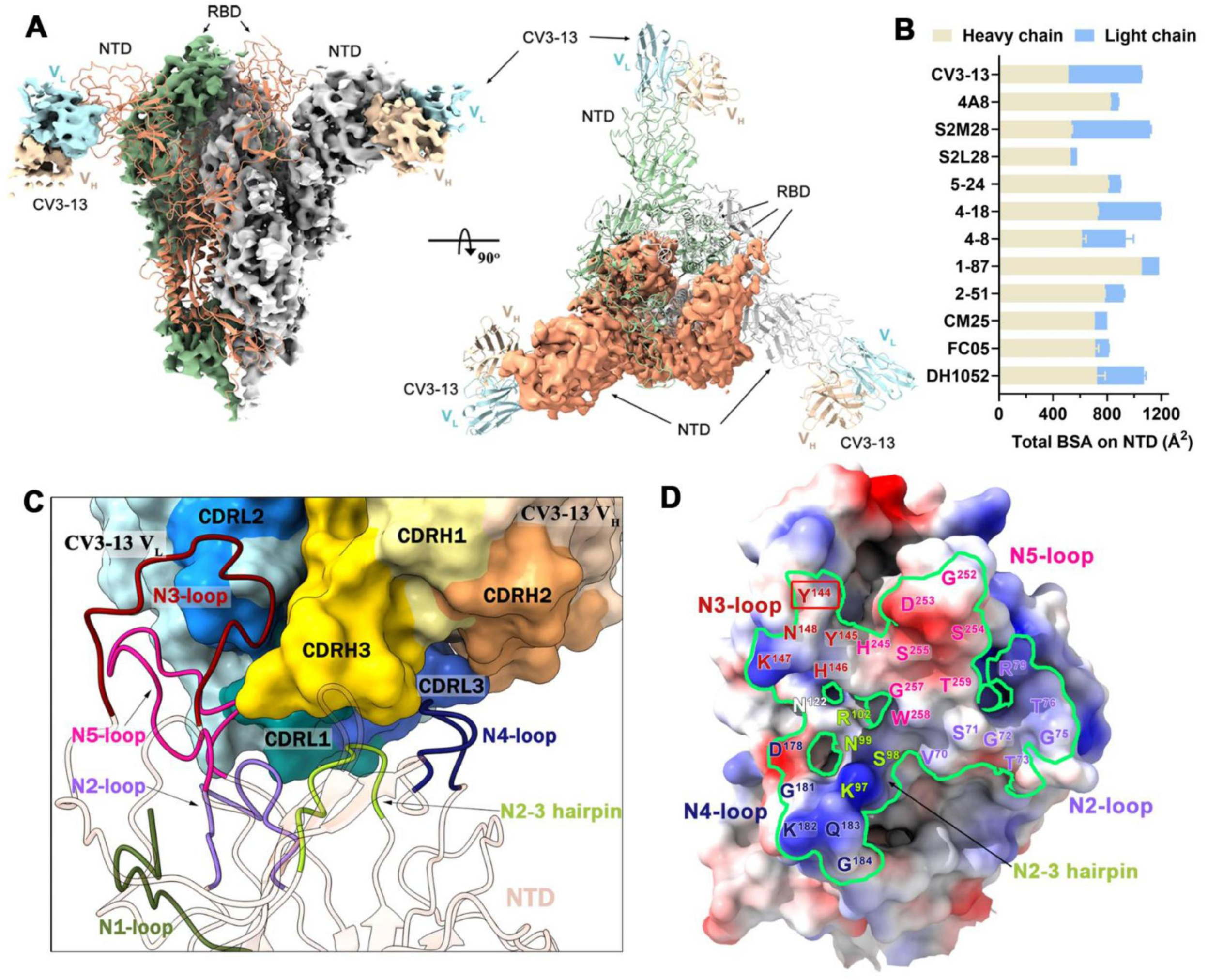
Cryo-EM structure of SARS-CoV-2 Spike in complex with CV3-13 Fab. (A) Side and top views of the NTD-targeting CV3-13 Fab and SARS-CoV-2 Spike complex reveal a symmetrical Fab-spike assembly. In the left panel, protomer A of the Spike colored in salmon is shown in ribbons while the other two protomers colored in green or grey and the variable regions (heavy chain - light yellow and light chain - cyan) of three CV3-13 Fabs binding to the lateral surface of the NTD are shown as cryo-EM density (C3-symmetry map). (B) The total buried surface area (BSA) at the Fab/NTD interface contributed by heavy/light chains of CV3-13 and eleven other structurally available NTD-directed mAbs,:4A8 (PDB: 7CL2), S2M28 (7LY3), S2L28 (7LXX), 5-24 (7L2F), 4-18 (7L2E), 4-8 (7LQV), 1-87 (7L2D), 2-51 (7L2C), CM25 (7M8J), FC05 (7CWS) and DH1052 (7LAB). The BSA values of all the equivalent biological assemblies were calculated and averaged by PISA (Krissinel and Henrick, 2007). (C) Expanded view of CV3-13 interactions with the protruding NTD loops. CV3-13 is shown as surface with the CDRs H1, H2, H3 L1, L2 and L3 colored in yellow, orange, gold, dark cyan, sky blue and deep blue respectively. The NTD regions are displayed as ribbons and the N1 to N5 loops as defined by (Chi et al., 2020) are colored in green, purple, deep red, blue and pink respectively. Of note, the newly identified N2-3 hairpin is highlighted in light green. (D) CV3-13 epitope footprint on the electrostatic potential surface of NTD (colored red, blue and white for negative, positive and neutral electrostatic potential respectively). The NTD residues interacting with CV3-13, which were defined as those with BSA > 0 Å^2^ as calculated by PISA, are colored as in (C) in accordance with loop locations. The CV3-13 epitope footprints is outlined in green. The deleted Y^144^ identified in B.1.1.7 variant that disrupts CV3-13 binding is marked with a red box.

The structure of CV3-13 Fab bound to SARS-CoV-2 HexaPro Spike is shown in Figure 3A. The variable heavy and light chain regions (V_H_ and V_L_) of CV3-13 were well defined in the density. The constant part of the Fab was disordered, however, and is omitted from the model. CV3-13 binds to the NTD, at an almost right angle relative to the stem region of the spike. Both heavy and light chain complementarity determining regions (CDRs) of CV3-13 contribute almost equally to antigen recognition with CDR H3, CDR L1 and CDR L3 having the greatest number of contacts. The total buried surface area (BSA) in complex formation is 1184 Å^2^ which is comparable to the BSA of typical complexes formed by other NTD-specific antibodies (Figure 3B). Four out of five highly mobile NTD loops as defined by Chi et al., (Chi et al., 2020), e.g.: N2 (residues 67-79), N3 (residues 141-156), N4 (residues 177-186) and N5 (residues 246-260) are stabilized by the associated CV3-13 Fab (Figure 3C). The fifth antigenic NTD loop N1 (residues 14-26) is distal from the CV3-13 binding site and exhibits a higher degree of mobility in the complex. Of the antibody CDRs, the 14-amino-acid CDR H3, consisting of 7 aromatic residues (six tyrosines and one phenylalanine), stretches into the hydrophilic groove formed by the positively charged N3 and N4 loops, the N2 and N3 loops, and the glycan- shielded Asn^122^. CV3-13 forms extensive contact with N2, N4 and a buried N2 to N3 loop hairpin (residues 95-102) which are rarely involved in the epitopes of other known NTD-mAbs (Figure 4C). Interestingly, CV3-13 relies on key π-π interactions through its CDR H3 tyrosines to the Tyr^144^-Tyr^145^ stretch of N3 that is frequently mutated in SARS-CoV-2 emerging variants. For example, the single residue deletion of Tyr^144^ (as first detected in the B.1.1.7 variant) which was used to initially characterize CV3-13 causes a complete abrogation of CV3-13 binding (Figure 1B).

**Figure 4.**
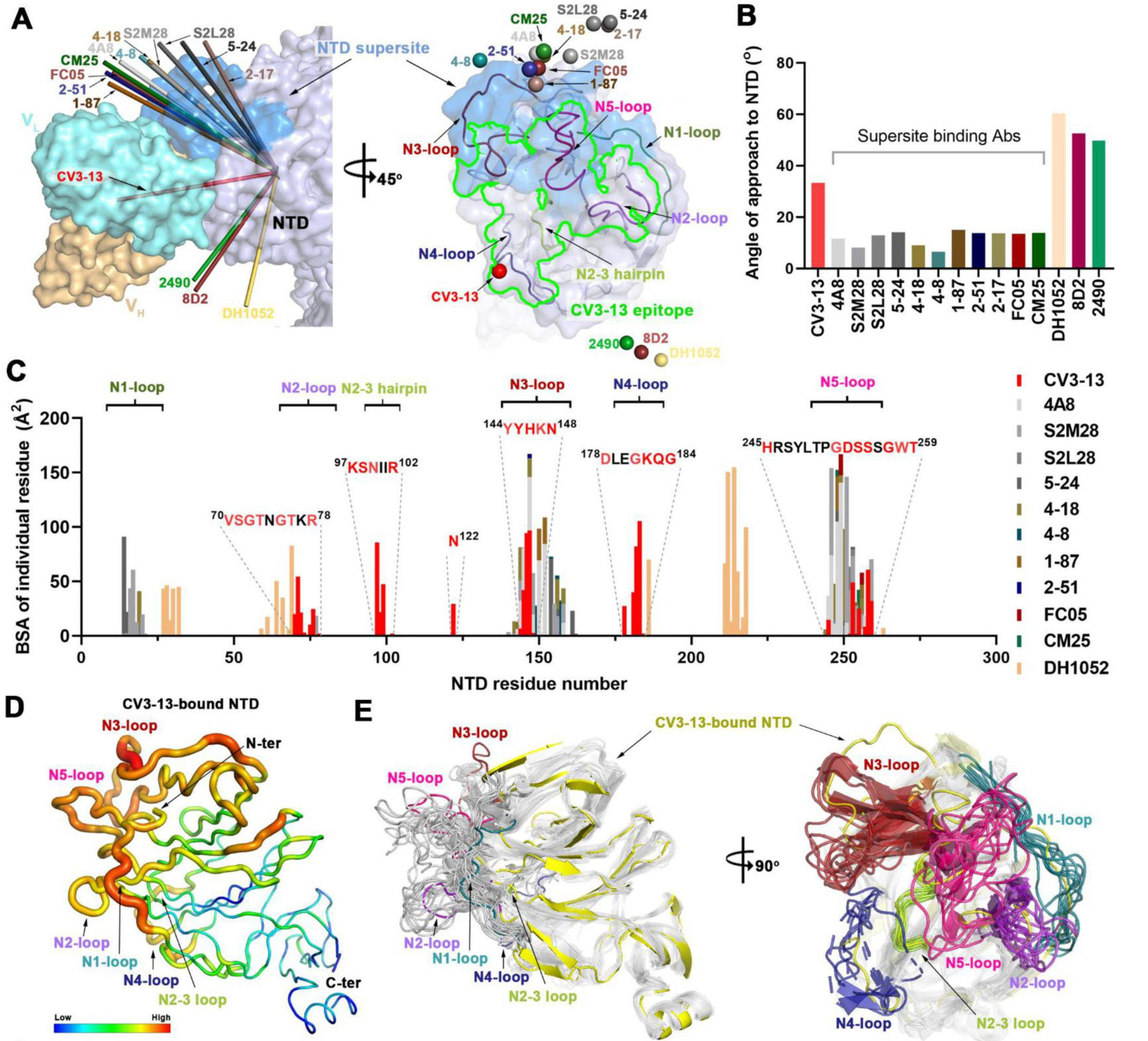
Structural basis of SARS-CoV-2 Spike recognition by CV3-13 and NTD- directed neutralizing mAbs. (A) Two-view diagram of CV3-13 binding to the NTD lateral epitope as compared to 11 other NTD-targeting neutralizing mAbs and 3 other infectivity enhancing mAbs. The NTD defined in this study is used here and shown as a light blue surface with the NTD supersite (residues 14-20, 140-158 and 245-264) highlighted in blue. On the left panel, the CV3-13 variable region is shown as a surface in light yellow for heavy chain and cyan for light chain. The approach of other NTD-binding mAbs are graphically represented by arrows (using the average C α position of the variable region for individual NTD-targeting antibodies (Cα-_Fv_) pointing toward the average Cα position of the NTD domains as a whole (Cα-_NTD_) determined by individual NTD/spike-mAb complexes) in indicated colors. On the right panel, the NTD is shown in semi-transparent surface with NTD loops N1-N5 and the newly defined N2-3 hairpin shown as colored ribbons. The CV3-13 epitope boundary is shown with a green line. The Cα-Fv of each NTD antibody is shown as colored spheres. (B) Angles of approach of CV3-13 and other NTD antibodies. The values for individual mAbs are calculated as the angle between Cα- _Fv_ and Cα-_NTD_ with Cα_-NTD-Cter_ acting as the origin. (C) Diagram of NTD-antibody epitopes reveal a distinct set of CV3-13 interacting NTD residues. The buried surface area (BSA) for the NTD residues contacting individual mAbs were calculated by PISA. (D) B-factor representation of the CV3-13-bound NTD domain. (E) Orthogonal views of the NTD superimposition of reported EM/X-ray structures of NTD-binding mAbs in complex with SARS-CoV-2 spike or NTD. On the left panel, N1-N5 loops and N2-3 hairpin of CV3-13-bound NTD are shown as colored ribbons with the NTD scaffold is colored in yellow, while the other NTDs are uniformly depicted as grey ribbons. In the right panel, CV3-13-bound NTD is shown by yellow ribbons and the antigen NTD loops from other mAb-NTD structures are shown with colored ribbons with the scaffold colored in grey.

The footprint of CV3-13 epitope differs from all known NAbs that recognize the NTD. Neutralizing NTD-directed antibodies target similar glycan-free epitopes located in the upper protruding area of the NTD, referred to as the “NTD supersite” of vulnerability (Cerutti et al., 2021; Harvey et al., 2021; McCallum et al., 2021). The non-neutralizing CV3-13 targets a distinct lateral region of the spike closer to the viral membrane surface, which only marginally overlaps with the supersite (Figure 4A). The distinct angle of approach of CV3-13 enables its extended CDR H3 to access the buried N2 to N3 hairpin which does not interact with the reported NTD-directed mAbs. Given that the NTD mutations found in escape variants of SARS-CoV-2 are thought to be the direct result of NTD-specific antibody selection, the emergence of the T95I mutation in the N2 to N3 hairpin, as first seen in the B.1.526 (Iota) and B.1.617.1 (Kappa) variants (Figure S6), suggests that this region represents a new antigenic site that can be targeted by non- neutralizing antibodies. Interestingly, the immunogenic N3 and N5 loops, which form the major interacting motifs for currently characterized NTD mAbs and is the primary component of the NTD supersite, only makes minor contact with the CDR H3, L1 and L2 of CV3-13. As a result, the backbone residues of the N3 and N5 loops are less traceable in the density map (Figure S5) and appear to adopt markedly different conformations as compared with those observed for neutralizing anti-NTD mAb structures (Figure 4E). Overall, the N3 and N5 loops which form the majority of the NTD supersite are less important for CV3-13 binding. Instead, CV3-13 utilizes the N2 and N4 loops and N2 to N3 hairpin, which are located at the lateral/bottom side of the NTD and are rarely accessed by the NTD-directed NAbs that bind from the top of the NTD. Our data are consistent with the finding that N3 and N5 loop engagement is an important component of NTD directed antibody neutralization of SARS-CoV-2.

### CV3-13 has a distinct angle of approach and induces conformational rearrangements in the NTD as compared to other NTD-binding mAbs

To gain structural insight on how the antibody recognition site and its angle of approach to the SARS-CoV-2 spike NTD affects the mode of action (i.e., potent neutralizers targeting the supersite versus non-neutralizing antibodies), we aligned the NTD-Fv portions of CV3-13 and other reported NTD-directed antibodies based on the rigid NTD core. The comparisons consisted of antigen-Fab structures for whose PDB models are available. There are in total eleven neutralizing antibodies targeting the NTD supersite as well as the other three which recognize the so-called infectivity-enhancing site (Li et al., 2021a; Liu et al., 2021; McCallum et al., 2021). As shown in Figure 4A, CV3-13 approaches the NTD at a nearly perpendicular angle relative to the spike trimer axis with its epitope footprint only partly overlapping the NTD supersite and the infectivity-enhancing site. The supersite binding antibodies access the NTD from the top of the spike trimer, while antibodies targeting the infectivity-enhancing site approach the spike from the bottom, closer to the viral membrane. The differences in binding modes of CV3-13 and other NTD-specific antibodies are evident when the angles of approach (defined as the angle between the average Cα position for the Fv of each individual mAb (Cα-Fv), and the average Cα position for the NTD as a whole (Cα-NTD), using the average Cα position of the C-terminal helix (residues 295-303) of the NTD (Cα-NTD-C-term) as the origin to exclude any differences due to the conformational state of the NTD relative to the spike as a whole) are calculated (Figure 4B). The supersite-binding NAbs approach the NTD with a similar angle (in a range of 6° -15°), substantially different from that of CV3-13 with a calculated angle ∼30° (Figure 4B). In contrast, the infectivity-enhancing antibodies use the angle of approach in the range of 45°-60°. Also, the fine epitope specificity of CV3-13 is different with contact regions only partially overlapping with neutralization supersite (Figure 4C). To summarize, CV3-13 uses the binding angle that positions it somewhere between the binding angles of antibodies recognizing the neutralization supersite, that bind from the top of the spike and antibodies targeting the infectivity-enhancing site, that bind at the bottom of spike, closer to the viral membrane. These features are likely why CV3-13 lacks direct neutralizing activity but has no infectivity-enhancing properties (Figure 2A, B). The CV3-13 binding mode permits effective engagement of innate immune cells to mediate Fc-effector activity. Indeed, it has been shown that antibodies targeting the NTD supersite have largely overlapping epitope footprints (all engaging the N1, N3 and N5 NTD loops) and a narrow range in their angle of approach that could allow them to sterically disrupt spike-receptor interactions, TMPRSS2-dependent activation and/or viral-host membrane fusion.

Both CV3-13 and the supersite specific Abs bind to the NTD utilizing highly mobile and conformationally unconstrained NTD loops by an induced-fit mechanism. To assess the impact of antibody recognition on the conformation of NTD, we superimposed the NTD domain from the ligand-free HexaPro Spike (PDB:6XKL) and eleven NTD-directed antibody-spike/NTD complexes and examined mAb-induced structural rearrangements. As described earlier by Cerutti and colleagues (Cerutti et al., 2021), the highly mobile N1-N5 loops (Figure 4D), which are largely disordered in the ligand-free spike, have diverse conformations in response to the bound Fabs at different sites (Figure 4E). The degree of local flexibility for the NTD loops is inversely correlated with their contribution to the Fab-NTD interface. These structural changes are a direct consequence of antibody binding, either by a conformational sampling or an induced-fit mechanism. Overall, NTD-directed antibodies induced substantial structural rearrangement of the NTD. The high immunogenicity and high mutational frequency observed in the flexible NTD loops likely results from their increased accessibility on the Spike and their subsequent encounters with both neutralizing and non-neutralizing antibodies.

### CV3-13 does not protect K18-hACE2 mice from a SARS-CoV-2 lethal challenge

We next investigated the *in vivo* efficacy of the non-neutralizing antibody CV3-13 under a prophylactic or therapeutic setting to protect or treat K18-hACE2 transgenic mice from a lethal SARS-CoV-2 infection. CV3-13 was delivered intraperitoneally (ip, 12.5 mg IgG/kg body weight) 24 h before (prophylactic) or one and two days post infection (dpi, therapeutic) in K18-hACE2 mice intranasally challenged with SARS-CoV-2 nLuc, as previously described (Figure S2 and S3) (Ullah et al., 2021). Longitudinal non-invasive bioluminescence imaging (BLI), body weight change, survival, viral load estimation in brain, lung and nasal cavity and terminal imaging after necropsy showed no difference between isotype- and CV3-13-treated mice under either prophylactic or therapeutic settings (Figure S2 and S3, panels A-J). Furthermore, mRNA levels of inflammatory cytokines (*IL6, CCL2, CXCL10* and *IFNG*) in brain and lungs were also similar to isotype-treated cohorts (Figure S2 and S3, panels K, L). These data suggest that natural Fc-effector functions associated with CV3-13 are not enough to alter virus replication and dissemination *in vivo* in the stringent K18-hACE2 mouse model of SARS-CoV-2 infection. Thus, in addition to clearing infected cells by engaging innate immune cells through Fc, efficient Fab-mediated neutralization of free viruses is essential for optimal *in vivo* efficacy. A compromise in either function reduces antibody-mediated protection *in vivo*.

### Fc-effector function enhancing mutations in CV3-13 delay neuroinvasion and mortality in a prophylactic regimen in K18-hACE2 mice

We explored if introducing Fc-effector function enhancing mutations (GASDALIE) in CV3-13 can improve *in vivo* efficacy against SARS-CoV-2. As shown in Figure 2, introduction of the GASDALIE mutations significantly enhanced Fc-mediated effector functions without affecting the lack of neutralizing activity of CV3-13. We carried out temporal BLI to visualize the viral dissemination profile when a CV3-13 GASDALIE mutant was administered under the prophylactic regimen in SARS-CoV-2-nLuc challenged K18-hACE2 mice (Figure 5A). Non-invasive imaging analyses revealed that CV3-13 GASDALIE-pre-treated mice showed reduced viral spread and delayed neuroinvasion compared to isotype-treated controls (Figure 5B, D, E). Accordingly, CV3-13 GASDALIE-treated mice displayed a decelerated body-weight loss phenotype and a significant delay in mortality (Figure 5F, G). Imaging animals at the experimental endpoint after necropsy revealed that systemic virus spread as measured by flux as well as viral loads in the nose, brain and lungs were significantly reduced in CV3-13 GASDALIE-treated compared to isotype-treated mice (Figure 5B, C, H-J). While the overall inflammatory cytokine mRNA expression profile in the brain and lungs were not different, IFN-gamma mRNA levels were significantly reduced in the brain tissues of CV3-13 GASDALIE-treated mice as compared to controls.

**Figure 5.**
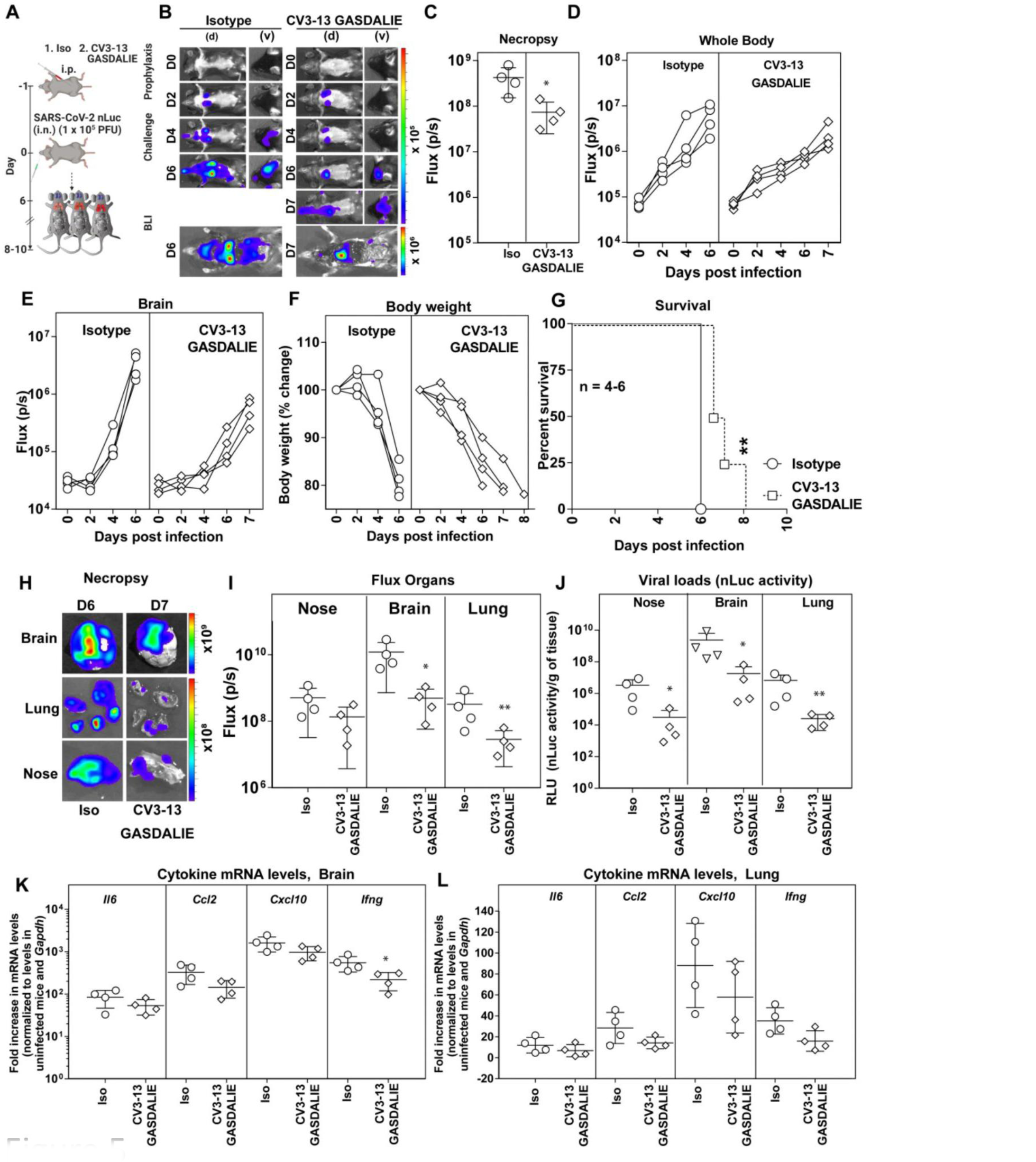
Prophylactic treatment with CV3-13 GASDALIE reduces SARS-CoV-2 spread and delays mortality in K18-hACE2 mouse model. (A) Experimental design for testing *in vivo* efficacy of Fc-effector function enhanced CV3-13 GASDALIE 1 day prior to challenging K18-hACE2 mice (i.n.) with SARS- CoV-2-nLuc followed by non-invasive BLI every 2 days. Human IgG1-treated (12.5 mg IgG/kg) mice were use as the isotype control (Iso) (B) Representative images from BLI of SARS-CoV-2-nLuc-infected mice in ventral (v) and dorsal (d) positions at the indicated dpi and after necropsy for experiment as in A. (C) *Ex-vivo* quantification of nLuc signal as flux (photons/sec) after necropsy. (D, E) Temporal quantification of nLuc signal as flux (photons/sec) computed non-invasively in indicated areas of each animal. (F) Temporal changes in mouse body weight with initial body weight set to 100 %. (G) Kaplan-Meier survival curves of mice statistically compared by log-rank (Mantel-Cox) test for experiment as in A. (H, I) *Ex-vivo* imaging of indicated organs and quantification of nLuc signal as flux (photons/sec) at the indicated dpi after necropsy. (J) Viral loads (nLuc activity/g) from indicated organs using Vero E6 cells as targets. (K, L) Cytokine mRNA levels in lung and brain tissues after necropsy normalized to *Gapdh* in the same sample and that in uninfected mice. Viral loads (J) and inflammatory cytokine profile (K,L) were determined after necropsy for mice when they succumbed to infection. Scale bars in (B) and (H) denote radiance (photons/sec/cm^2^/steradian). Each curve in (D)-(F) and each data point in (C, I)-(L) represents an individual mouse. The data in (C), (I)-(L) were analyzed by Mann Whitney U test. Statistical significance to isotype control are shown. ∗, p < 0.05; ∗∗, p < 0.01;1; Mean values ± SD are depicted.

To confirm the ability of CV3-13 GASDALIE to reduce virus spread and delay neuroinvasion, we terminated the experiment at 4 dpi (Figure 6A). Our BLI analyses confirmed a significant decrease in nLuc signals in the brain both non-invasively and after necropsy (Figure 6B-E, G, H). Viral burden measurements by assessing Nucleocapsid mRNA levels and nLuc activity from Vero ACE2-TMPRSS2 infected with tissue homogenates as well as histological analyses corroborated a significant reduction of viral replication in the brains of CV3-13 GASDALIE-treated mice (Figure 6B-E, G- K). While viral loads and cytokine mRNA levels in the lungs were similar between the two cohorts, there was a significant decrease in overall inflammatory cytokine mRNA levels in the brain consistent with reduced viral loads in CV3-13 GASDALIE-treated mice (Figure 6L, M). Thus, these data demonstrate the potential of non-neutralizing antibodies with enhanced Fc-effector functions to interfere with SARS-CoV-2 spread.

**Figure 6.**
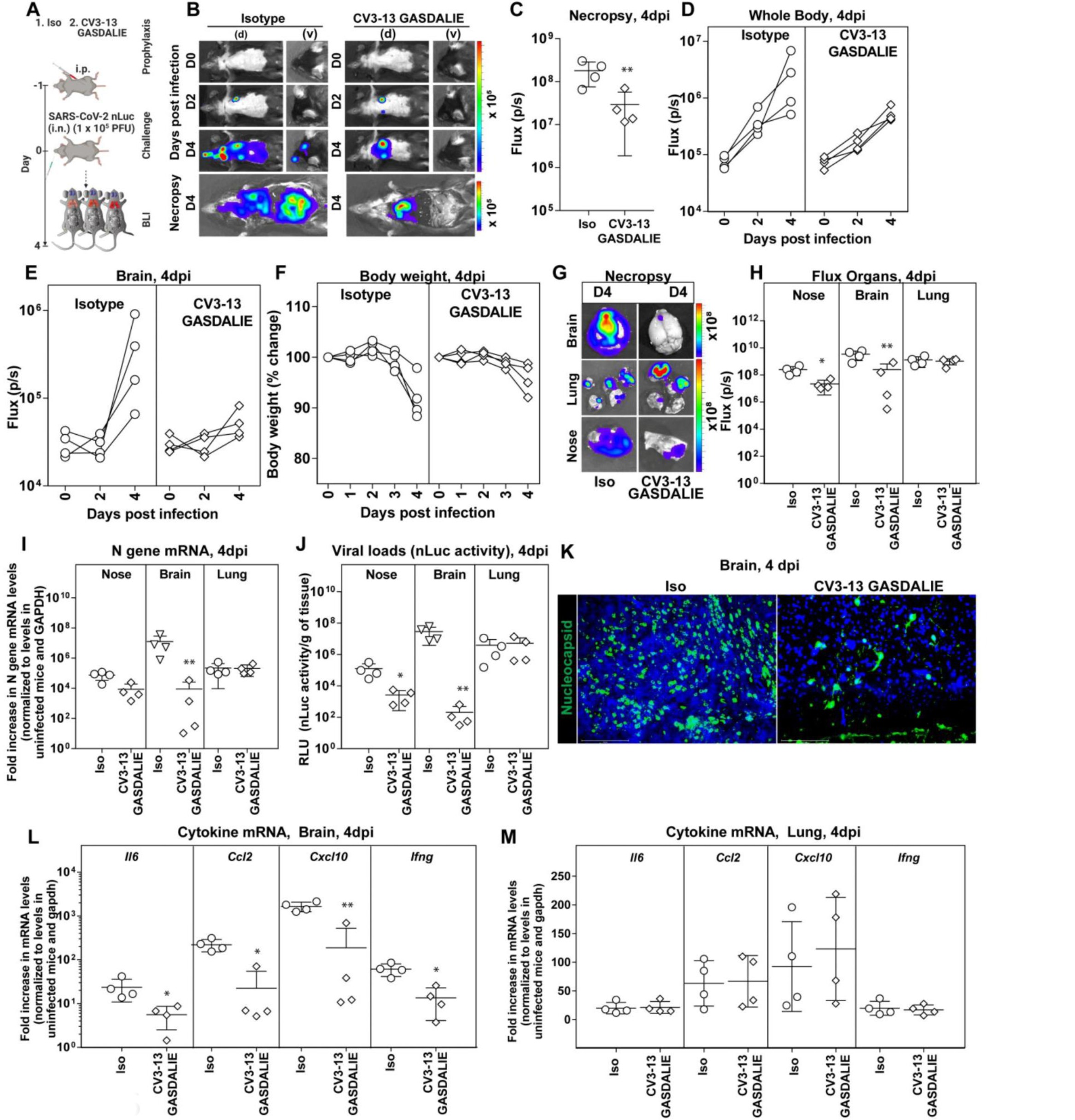
CV3-13 GASDALIE delays neuroinvasion and reduces inflammation in the brain of SARS-CoV-2-challenged K18-hACE2 mice. (A) Experimental design for testing virus dissemination in CV3-13 GASDALIE administered K18-hACE2 mice, 1 day prior to challenging (i.n.) with SARS-CoV-2-nLuc followed by non-invasive BLI every 2 days. Human IgG1-treated (12.5 mg IgG/kg) mice were use as the isotype control (Iso). (B) Representative images from BLI of SARS-CoV- 2-nLuc-infected mice in ventral (v) and dorsal (d) positions at the indicated dpi and after necropsy at indicated days for experiment as in A. (C) *Ex-vivo* quantification of nLuc signal as flux (photons/sec) at 4 dpi after necropsy. (D, E) Temporal quantification of nLuc signal as flux (photons/sec) computed non-invasively in indicated areas of each animal. (F) Temporal changes in mouse body weight with initial body weight set to 100%. (G, H) *Ex-vivo* imaging of organs and quantification of nLuc signal as flux (photons/sec) at the indicated dpi after necropsy. (I) A plot showing real-time PCR analyses to detect SARS-CoV-2 nucleocapsid (N) gene mRNA in indicated organs at 4 dpi. The data were normalized to N RNA seen in uninfected mice and GAPDH mRNA levels. (J) Viral loads (nLuc activity/g) from indicated organs using Vero E6 cells as targets. (K) Images of cryosections from brain tissues of SARS-CoV-2-nLuc infected K18-hACE2 mouse treated with indicated antibodies and harvested at 4 dpi. Actin (blue) was detected using phalloidin-CF450, nucleocapsid (green) was detected using anti- SARS-CoV-2 N conjugated to AF488. Scale bar 150 µm. (L, M) Cytokine mRNA levels in lung and brain tissues after necropsy normalized to *Gapdh* in the same sample and that in uninfected mice. Viral loads (J) and inflammatory cytokine profile (L, M) were determined after necropsy for mice that succumbed to infection. Scale bars in (B) and (G) denote radiance (photons/sec/cm^2^/steradian). Each curve in (D)-(F) and each data point in (C), (H)-(J), (L) and (M) represents an individual mouse. The data in (C), (H)-(J), (L) and (M) were analyzed by Mann Whitney U test. Statistical significance to isotype control are shown. ∗, p < 0.05; ∗∗, p < 0.01; Mean values ± SD are depicted.

## Discussion

Neutralization plays a crucial role in protection against SARS-CoV-2 infection. Therefore many studies have focused on neutralization responses from convalescent plasma (Anand et al., 2021; Beaudoin-Bussieres et al., 2020; Gasser et al., 2021; Long et al., 2020; Prevost et al., 2020; Robbiani et al., 2020), vaccine-elicited antibodies (AstraZeneca and Iqvia Pty, 2021; Baden et al., 2021; Bio and Pfizer, 2021; Janssen and Prevention, 2021; ModernaTx et al., 2022; Polack et al., 2020; Skowronski and De Serres, 2021; Tauzin et al., 2021) and cocktails of mAbs for use as therapeutics (Hurlburt et al., 2020; Jennewein et al., 2021; Ju et al., 2020; Li et al., 2021b; Liu et al., 2020; Schafer et al., 2021; Tian et al., 2020; Wu et al., 2020c; Yuan et al., 2020). However, antibodies are polyvalent molecules able to mediate several antiviral functions (Adeniji et al., 2021). Among these, their capacity to recognize antigens at the surface of viral particles or infected cells and to recruit effector cells is gaining attention for SARS-CoV- 2 infection (Brunet-Ratnasingham et al., 2021; Schafer et al., 2021; Tortorici et al., 2020; Ullah et al., 2021; Winkler et al., 2021; Zohar et al., 2020). Also, recent studies on vaccine-elicited humoral responses suggest that additional mechanisms, besides neutralization, could be playing a role in vaccine efficacy (Alter et al., 2021; Amanat et al., 2021; Stankov et al., 2021; Tauzin et al., 2021). To address whether a non- neutralizing antibody can afford any level of protection from SARS-CoV-2 infection, we isolated a non-neutralizing antibody, CV3-13, from a convalescent donor and assessed its impact on the virus. CV3-13 binds the NTD of the Spike with high affinity and defines a novel NTD epitope. As compared to other NTD-supersite-directed neutralizing mAbs which bind the NTD predominately through use of the N1, N3 and N5 loops and approach the NTD from the top, CV3-13 targets the lateral NTD surface, engaging and rearranging a new set of antigenic NTD loops: N2, N4 and N2-3. The NTD interface residues, as identified by the cryo-EM structure, harbor frequent mutations in several circulating SARS-CoV-2 variants, including B.1.1.7 (alpha), B.1.617.1 (kappa), and C.37 (lambda), defining the structural basis for viral escape from CV3-13 and CV3-13-like antibodies. For instance, Tyr^144^ on the protruding N3 loop, forms close contact with the Tyr/Phe-rich CDR H3 of CV3-13 (Figure 3D), and is deleted in B.1.1.7, which markedly reduces its binding to CV3-13 (Figure 1A). Other NTD mutations outside of the supersite are found within the CV3-13 epitope including an N2 deletion ^69^HisVal^70^ in B.1.1.7, a Gly^75^ to Val/Thr^76^ to Ile double mutation in C.37, and a recurrent Thr^95^ to Ile mutation in B.1.526 and B.1.617.1 (Figure S6A). This is in line with the interpretation that NTD mutations from the emerging variants may be a result of NTD-directed antibody selection, suggesting that non-neutralizing NTD antibodies like CV3-13 influence virus evolution through the Fc-mediated effector functions.

CV3-13 did not neutralize pseudoviral particles or authentic SARS-CoV-2 viruses but did mediate Fc effector functions against Spike-expressing cells. We suggest that differences in fine epitope specificity and the angle of approach used by CV3-13 as compared to neutralizing NTD-specific mAbs limit its ability to sterically hinder Spike-co- receptor/auxiliary receptor interactions, the prefusion-to-postfusion transition of Spike and/or membrane fusion as has been suggested as a neutralization mechanism for other NTD binding antibodies.

Our data demonstrates that an antibody devoid of neutralizing activity is able to reduce virus dissemination and delay death in mice from lethal SARS-CoV-2 challenge via its Fc-mediated effector functions. While wild-type CV3-13 IgG1 did not provide any protection, CV3-13 GASDALIE mutant delayed death in prophylactically treated mice. These data suggest that a threshold of Fc-mediated effector function activity was required to impede virus progression.

While non-neutralizing antibodies do not directly inactivate free viruses, they remain attractive candidates for eliminating infected cells through Fc engagement and reduce virus burden. Although the characteristics including binding efficiency and angle of approach for efficient elimination of infected cells by non-neutralizing antibodies remains to be elucidated, here we show that enhancing Fc mutations (GASDALIE) are a crucial step in improving in vivo efficacy.

Altogether, CV3-13 represents of a new class of non-neutralizing NTD-directed mAbs that can mediate Fc-effector functions both *in vivo* and *in vitro*. Our data indicates that in addition to neutralization, additional antibody properties including Fc-mediated effector functions contribute to limiting viral spread and aid in fighting SARS-CoV-2 infection.

## Supplemental information

6 Supplementary figures and 1 table.

## Author contributions

GBB, YC, PDU, MP & AF : Conceptualization, Experimental design, Interpretation, Manuscript preparation and Writing; GBB : Generation of the GASDALIE and LALA mutants, Figure generation, Data processing and Initial draft; IU: Animal experiments, BLI, Viral load analyses, Data processing and Figure generation; YC, EP, WT, MP & FZ: Cryo-EM data collection, processing and validation; YC, WT & MP: Structural analysis and Figure generation; PDU: Histological analyses and Figure generation; GBB, JP, MB, SYG, SD, RG, YC, AT, GG, LS, DV, DC, ATM, JR, MP, WM & AF: Isolation, generation and characterization of SARS-CoV-2 S mAb CV3-13; SYG, JP & DV : Generation of the individual B.1.1.7 mutations in the Spike; MR & HC: Provided authentic SARS-CoV-2 virus; PK, MP, PDU, WM & AF: Funding for the work. Every author has read, edited, and approved the final manuscript.

## Acknowledgements

The authors thank the CRCHUM BSL-3 and Flow Cytometry Platforms for technical assistance. We thank Dr. Stefan Pöhlmann and Dr. Markus Hoffmann (Georg-August University) for the plasmids coding for SARS-CoV-2 and SARS-CoV-1 S and Dr. M. Gordon Joyce (U.S. MHRP) for the CR3022 mAb. Antibodies CV3-1, CV3-13 and CV3-25 were produced using the pTT vector kindly provided by the National Research Council of Canada. Schematics for showing experimental design in figures were created with BioRender.com. This work was supported by le Ministère de l’Économie et de l’Innovation (MEI) du Québec, Programme de soutien aux organismes de recherche et d’innovation, the Fondation du CHUM, a CIHR foundation grant #352417, an Exceptional Fund COVID-19 from the Canada Foundation for Innovation (CFI) #41027, a CIHR operating grant Pandemic and Health Emergencies Research/Project #177958 and a CIHR stream 1 and 2 for SARS-CoV-2 Variant Research to A.F. Support was also provided by the Sentinelle COVID Quebec network led by the Laboratoire de Santé Publique du Quebec (LSPQ) in collaboration with Fonds de Recherche du Québec-Santé (FRQS) and Genome Canada – Génome Québec, and by the Ministère de la Santé et des Services Sociaux (MSSS) and MEI to A.F. and M.R. NIH grant R01AI163395 to W.M. A.F. is recipient of a Canada Research Chair on Retroviral Entry no. RCHS0235 950- 232424. G.B.B. is the recipient of CIHR and FRQS PhD fellowships and J.P. is the recipient of a CIHR PhD fellowship. RG is supported by a MITACS Accélération postdoctoral fellowship. The funders had no role in study design, data collection and analysis, decision to publish, or preparation of the manuscript. CryoEM data were collected at the Maryland Center for Advanced Molecular Analyses which is supported by MPOWER (The University of Maryland Strategic Partnership).

## Disclaimer

The views expressed in this presentation are those of the authors and do not reflect the official policy or position of the Uniformed Services University, the U.S. Army, the Department of Defense, or the U.S. Government.

## STAR Methods

### KEY RESOURCES TABLE

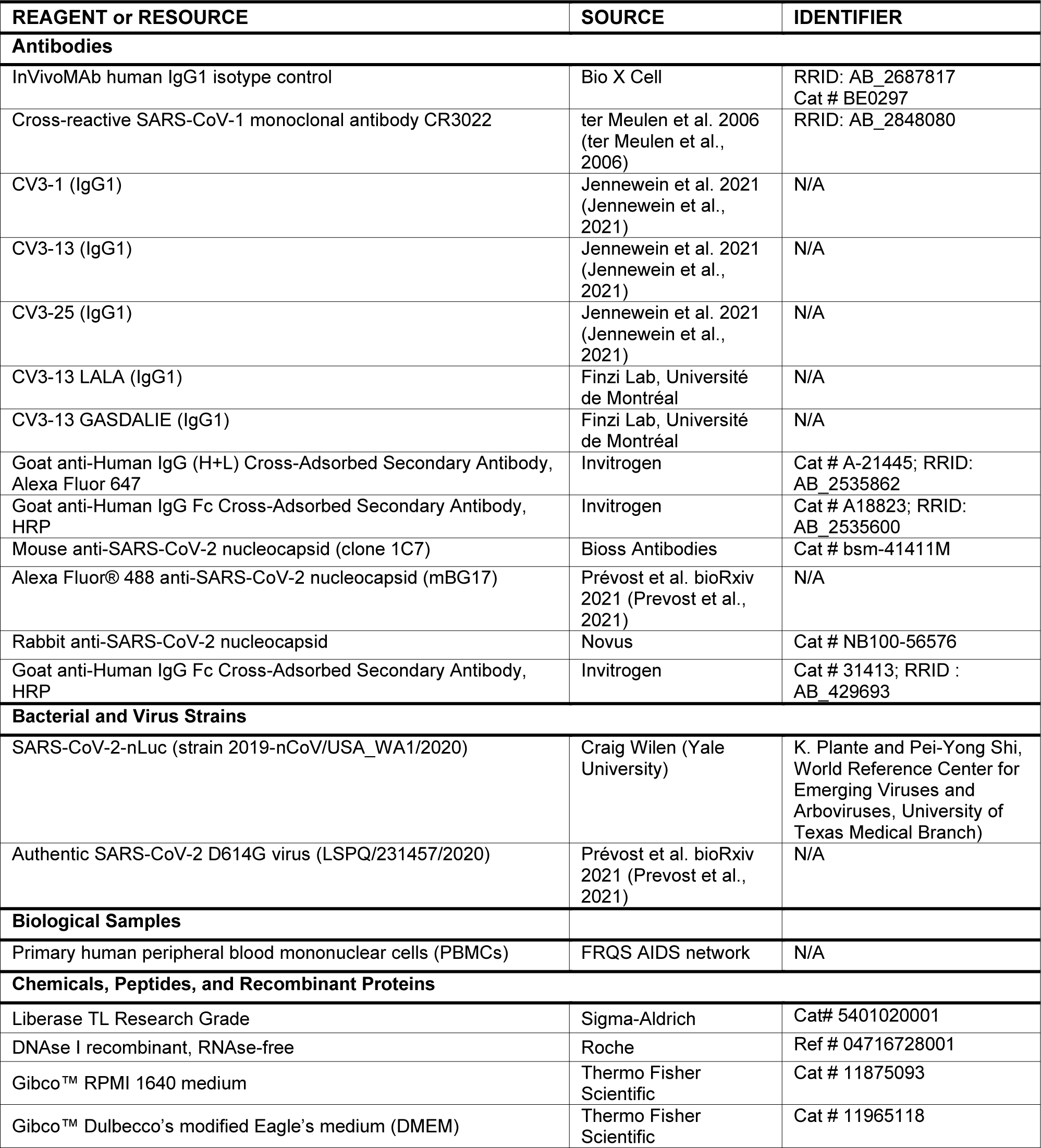
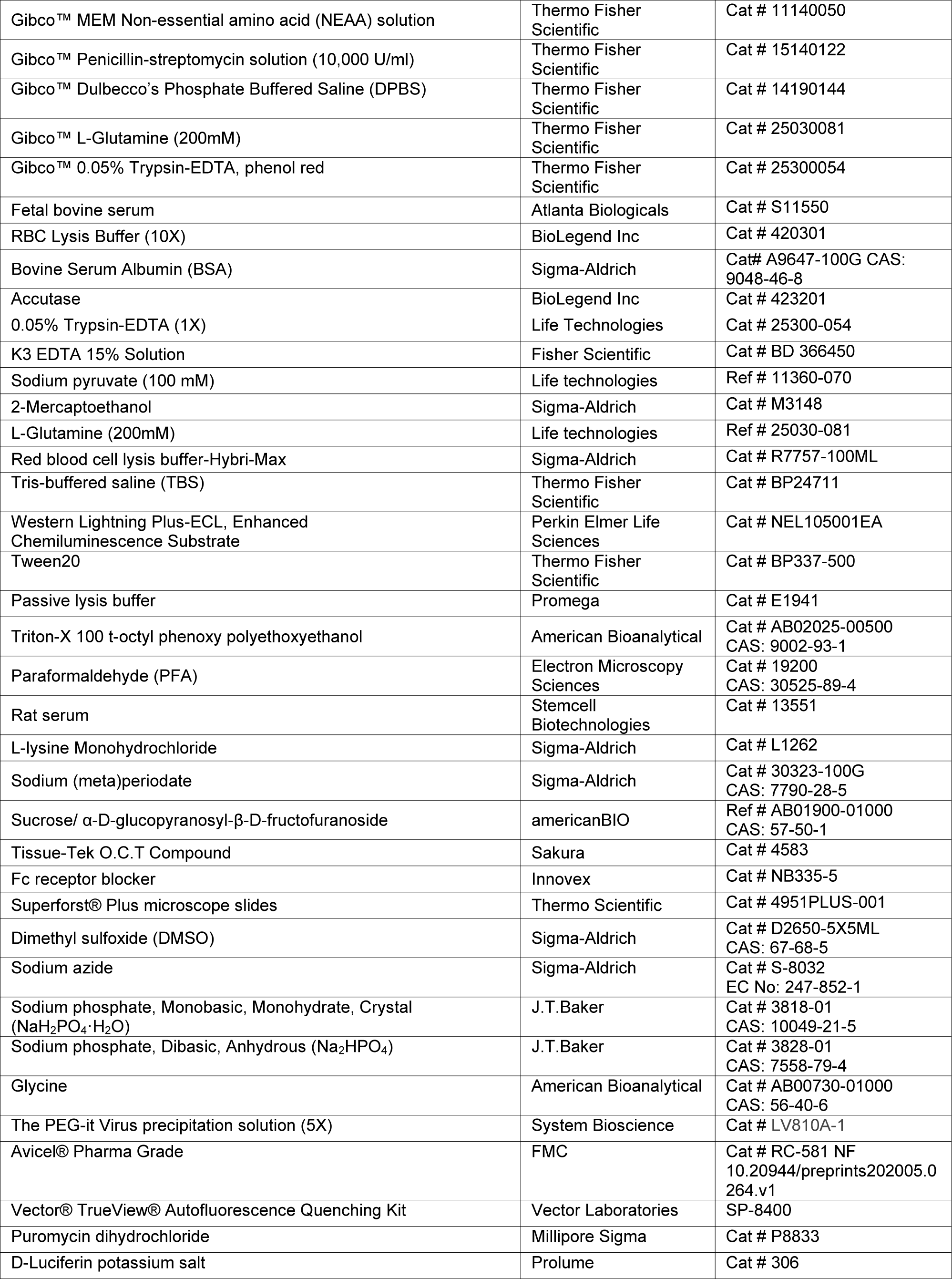
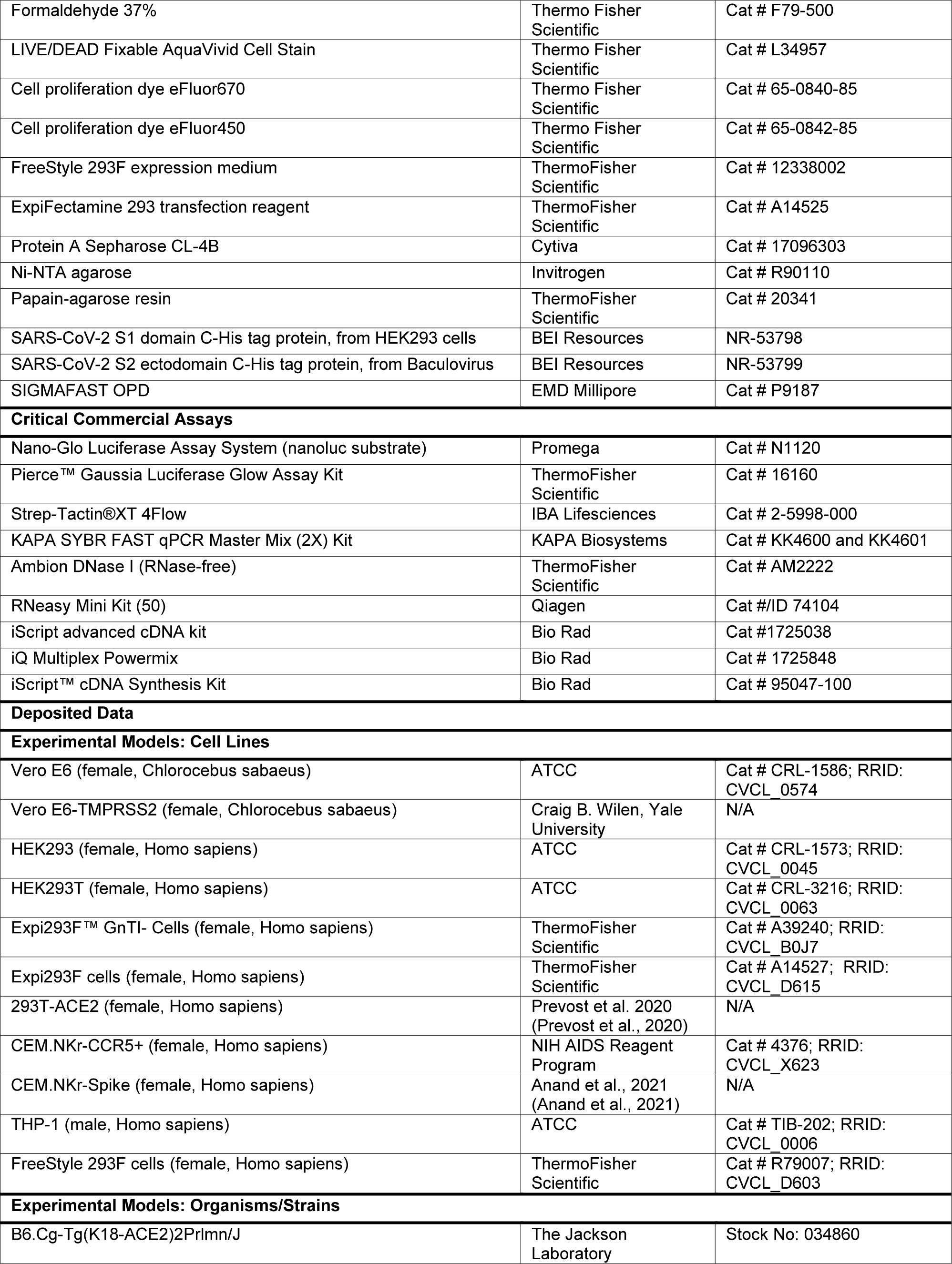
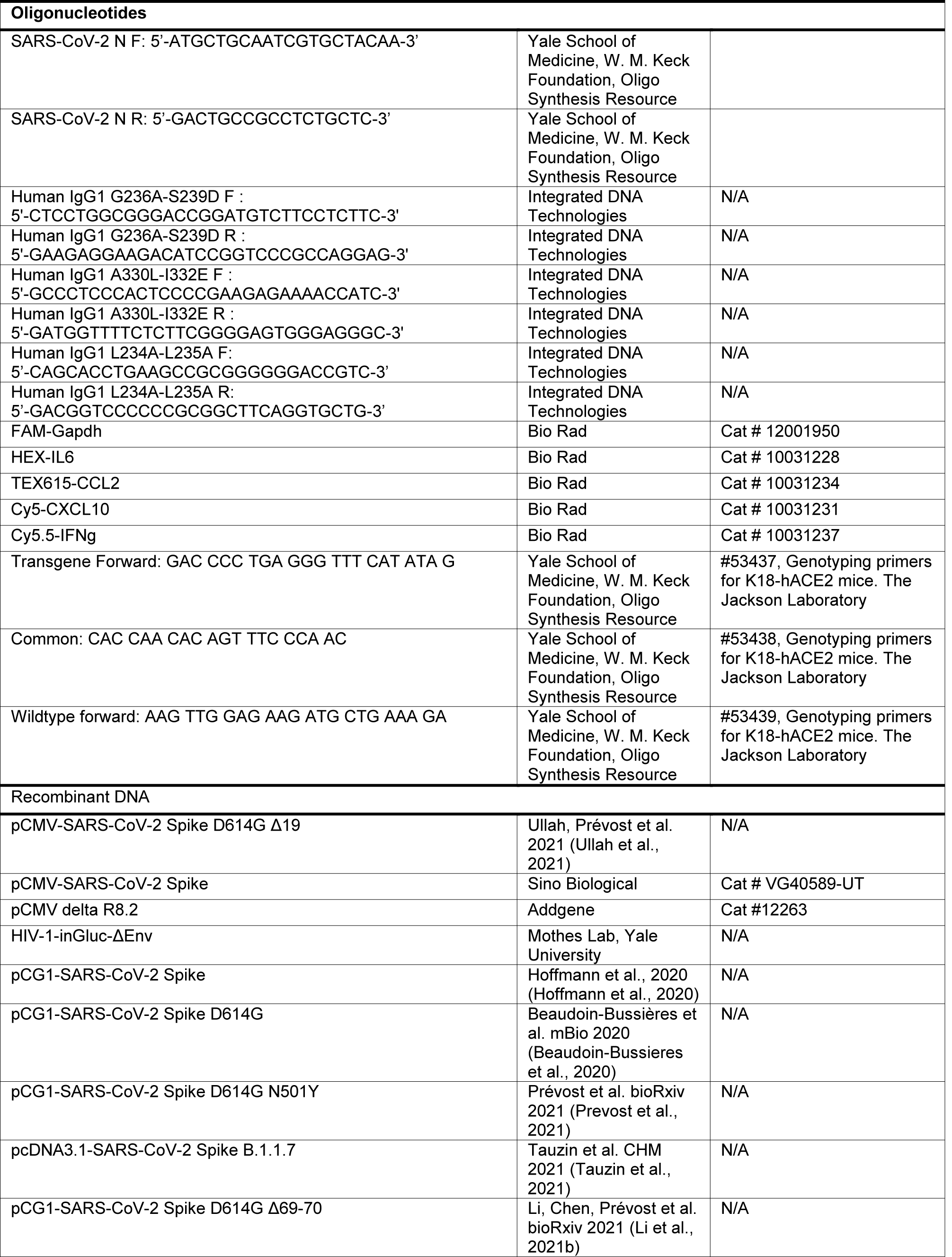
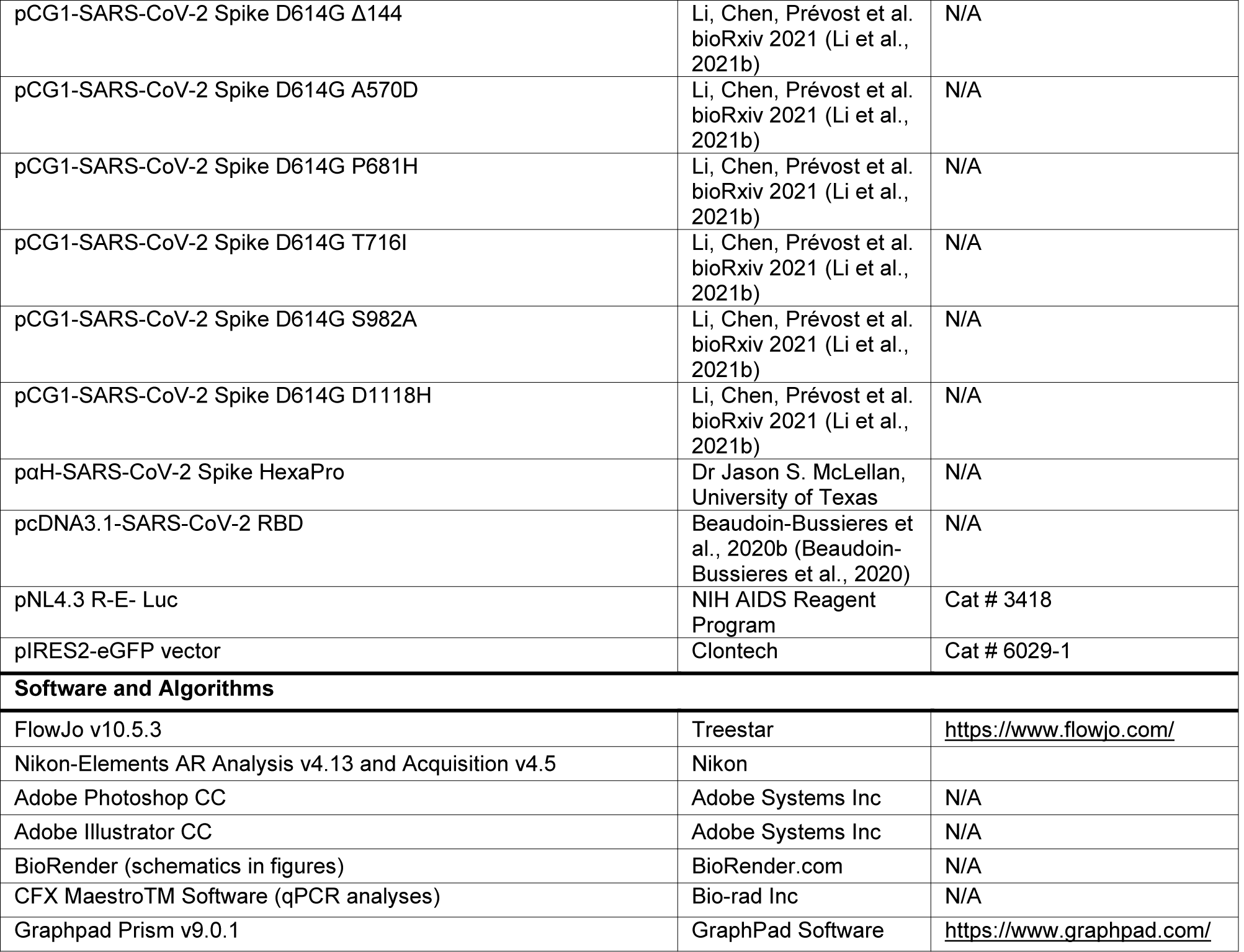
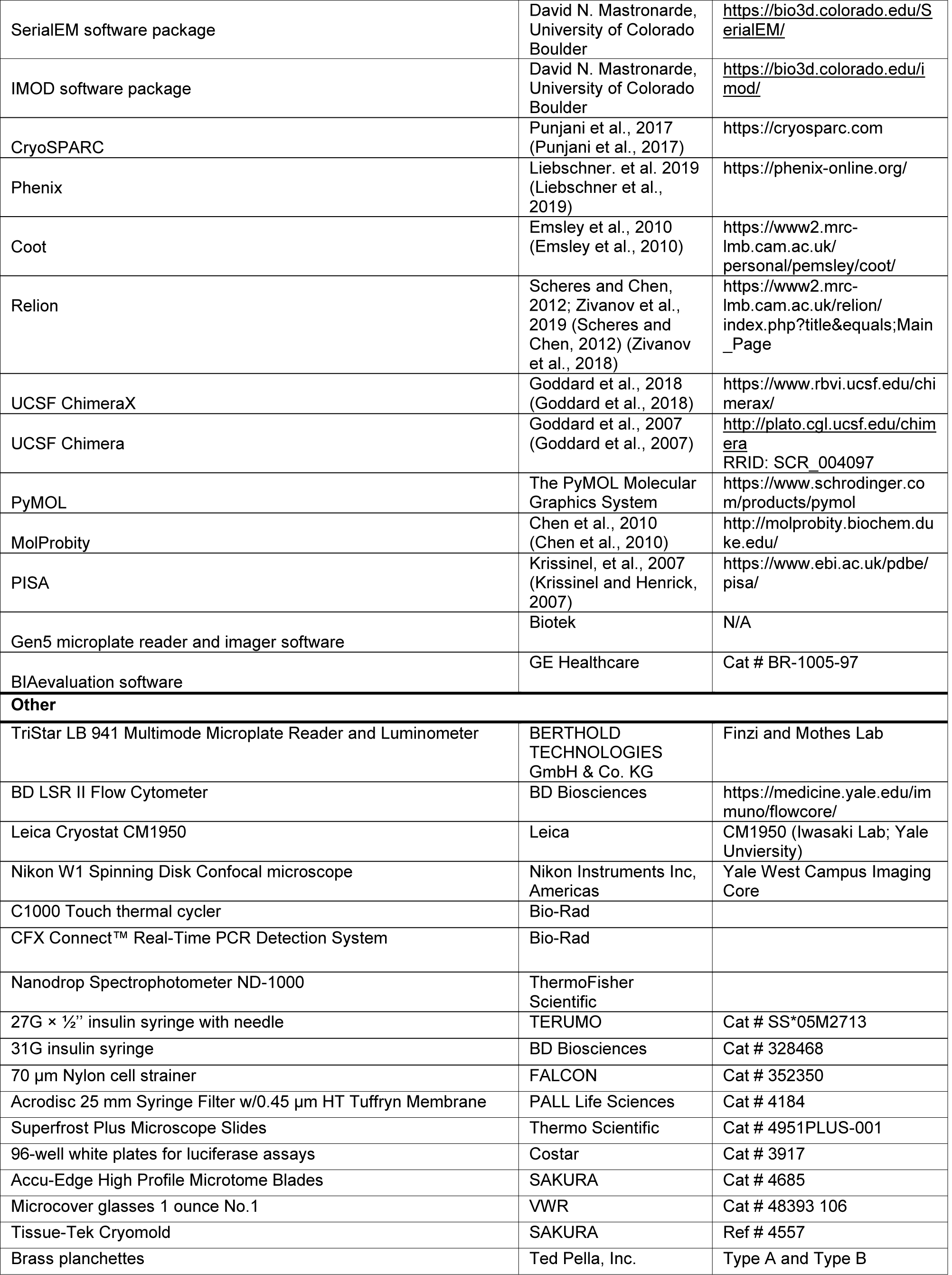
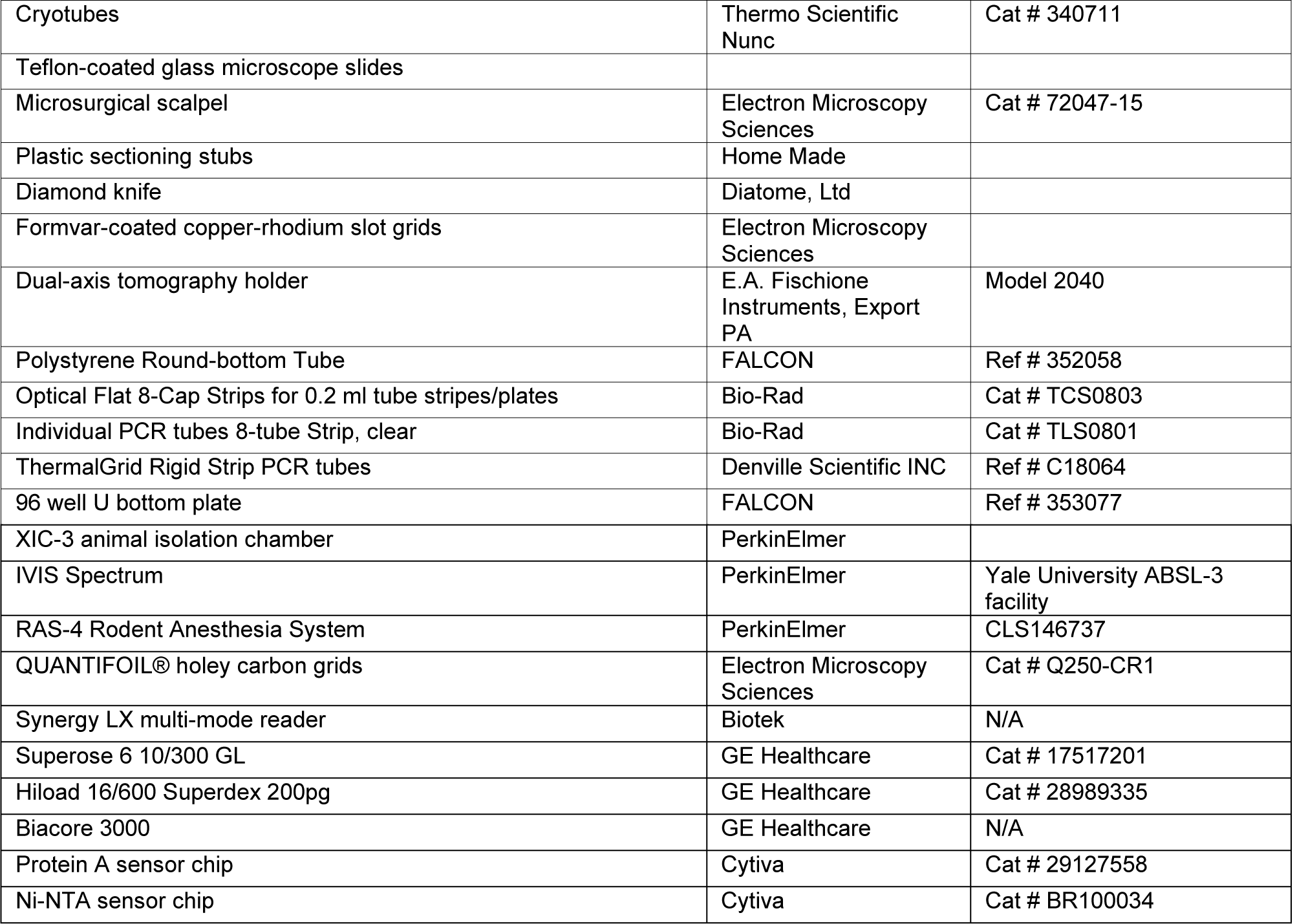

### RESOURCE AVAILABILITY

#### Lead Contact

Further information and requests for resources and reagents should be directed to and will be fulfilled by the Lead Contact, Andrés Finzi.

#### Materials Availability

All other unique reagents generated in this study are available from the corresponding authors with a completed Materials Transfer Agreement.

#### Data and Code Availability

The data that support the findings of this study are available from the corresponding authors upon reasonable request.

### EXPERIMENTAL MODEL AND SUBJECT DETAILS

#### Cell and Viruses

Vero E6 (CRL-1586, American Type Culture Collection (ATCC) or Vero E6-TMPRSS2 (Craig B. Wilen, Yale University), were cultured at 37°C in RPMI supplemented with 10% fetal bovine serum (FBS), 10 mM HEPES pH 7.3, 1 mM sodium pyruvate, 1× non-essential amino acids, and 100 U/ml of penicillin–streptomycin. The 2019n- CoV/USA_WA1/2019 isolate of SARS-CoV-2 expressing nanoluciferase was obtained from Craig Wilen, Yale University and generously provided by K. Plante and Pei-Yong Shi, World Reference Center for Emerging Viruses and Arboviruses, University of Texas Medical Branch) (Xie et al., 2020a; Xie et al., 2020b). Virus was propagated in Vero-E6 or Vero E6-TMPRSS2 by infecting them in T150 cm^2^ flasks at a MOI of 0.1. The culture supernatants were collected after 72 h when cytopathic effects were clearly visible. The cell debris was removed by centrifugation and filtered through 0.45-micron filter to generate virus stocks. Viruses were concentrated by adding one volume of cold (4 °C) 4x PEG-it Virus Precipitation Solution (40 % (w/v) PEG-8000 and 1.2 M NaCl; System Biosciences) to three volumes of virus-containing supernatant. The solution was mixed by inverting the tubes several times and then incubated at 4 °C overnight. The precipitated virus was harvested by centrifugation at 1,500 × g for 60 minutes at 4 °C. The concentrated virus was then resuspended in PBS then aliquoted for storage at −80°C. All work with infectious SARS-CoV-2 was performed in Institutional Biosafety Committee approved BSL3 and A-BSL3 facilities at Yale University School of Medicine or Université de Montréal using appropriate positive pressure air respirators and protective equipment. CEM.NKr, CEM.NKr-Spike, THP-1 and peripheral blood mononuclear cells (PBMCs) were maintained at 37°C under 5% CO_2_ in RPMI media, supplemented with 10% FBS and 100 U/mL penicillin/streptomycin. 293T (or HEK293T) and 293T-ACE2 cells were maintained at 37°C under 5% CO_2_ in DMEM media, supplemented with 5% FBS and 100 U/mL penicillin/streptomycin. CEM.NKr (NIH AIDS Reagent Program) is a T lymphocytic cell line resistant to NK cell-mediated lysis. CEM.NKr-Spike stably expressing SARS-CoV-2 Spike were used as target cells in ADCC and ADCP assays (Anand et al., 2021). THP-1 monocytic cell line (ATCC) was used as effector cells in the ADCP assay. PBMCs were obtained from healthy donor through leukapheresis and were used as effector cells in ADCC assay. 293T cells (obtained from ATCC) were derived from 293 cells, into which the simian virus 40 T-antigen was inserted. 293T-ACE2 cells stably expressing human ACE2 is derived from 293T cells (Prevost et al., 2020). 293T-ACE2 cells were cultured in medium supplemented with 2 mg/mL of puromycin (Millipore Sigma).

#### Ethics statement

PBMCs from healthy individuals as a source of effector cells in our ADCC assay were obtained under CRCHUM institutional review board (protocol #19.381). Research adhered to the standards indicated by the Declaration of Helsinki. All participants were adults and provided informed written consent prior to enrollment in accordance with Institutional Review Board approval.

#### Antibodies

The human antibodies (CV3-1, CV3-25 and CV3-13) used in the work were isolated from blood of male convalescent donor CV3 recovered 41 days after symptoms onset using fluorescent recombinant stabilized Spike ectodomains (S2P) as probes to identify antigen-specific B cells as previously described (Lu et al., 2020; Seydoux et al., 2020). Light and heavy chains were cloned into the pTT expression plasmid (Durocher et al., 2002). Site-directed mutagenesis was performed on plasmids expressing CV3-13 antibody heavy chains in order to introduce the LALA mutations (L234A/L235A) and the GASDALIE mutations (G236A/S239D/A330L/I332E) using to the QuickChange II XL site-directed mutagenesis protocol (Stratagene).

#### Mouse Experiments

All experiments were approved by the Institutional Animal Care and Use Committees (IACUC) of and Institutional Biosafety Committee of Yale University (IBSCYU). All the animals were housed under specific pathogen-free conditions in the facilities provided and supported by Yale Animal Resources Center (YARC). All IVIS imaging, blood draw and virus inoculation experiments were done under anesthesia using regulated flow of isoflurane:oxygen mix to minimize pain and discomfort to the animals.

### METHOD DETAILS

#### SARS-CoV-2 infection and treatment conditions

For all *in vivo* experiments, the 6 to 8 weeks male and female hACE2-K18 mice were intranasally challenged with 1 x 10^5^ PFU in 25-30 µl volume under anesthesia (0.5 - 5 % isoflurane delivered using precision Dräger vaporizer with oxygen flow rate of 1 L/min). For mAb treatment using prophylaxis regimen, mice were treated with 250 µg (12.5 µg/g body weight) of indicated antibodies (CV3-13 WT or CV3-13 GASDALIE) via intraperitoneal injection (i.p.) 24 h prior to infection. For mAb treatment under therapeutic regimen, mice were treated at 1 and 2 dpi intraperitoneally with CV3-13 (12.5 µg/g body weight). Body weight was measured and recorded daily. The starting body weight was set to 100 %. For survival experiments, mice were monitored every 6-12 h starting six days after virus administration. Lethargic and moribund mice or mice that had lost more than 20 % of their body weight were sacrificed and considered to have succumbed to infection for Kaplan-Meier survival plots.

#### Bioluminescence Imaging (BLI) of SARS-CoV-2 infection

All standard operating procedures and protocols for IVIS imaging of SARS-CoV-2 infected animals under ABSL-3 conditions were approved by IACUC, IBSCYU and YARC. All the imaging was carried out using IVIS Spectrum® (PerkinElmer) in XIC-3 animal isolation chamber (PerkinElmer) that provided biological isolation of anesthetized mice or individual organs during the imaging procedure. All mice were anesthetized via isoflurane inhalation (3 - 5 % isoflurane, oxygen flow rate of 1.5 L/min) prior and during BLI using the XGI-8 Gas Anesthesia System. Prior to imaging, 100 µL of nanoluciferase substrate, furimazine (NanoGlo^TM^, Promega, Madison, WI) diluted 1:40 in endotoxin-free PBS was retro-orbitally administered to mice under anesthesia. The mice were then placed into XIC-3 animal isolation chamber (PerkinElmer) pre-saturated with isothesia and oxygen mix. The mice were imaged in both dorsal and ventral position at indicated days post infection. The animals were then imaged again after euthanasia and necropsy by spreading additional 200 µL of substrate on to exposed intact organs. Infected areas of interest identified by carrying out whole-body imaging after necropsy were isolated, washed in PBS to remove residual blood and placed onto a clear plastic plate. Additional droplets of furimazine in PBS (1:40) were added to organs and soaked in substrate for 1-2 min before BLI.

Images were acquired and analyzed with the manufacturer’s Living Image v4.7.3 *in vivo* software package. Image acquisition exposures were set to auto, with imaging parameter preferences set in order of exposure time, binning, and f/stop, respectively. Images were acquired with luminescent f/stop of 2, photographic f/stop of 8. Binning was set to medium. Comparative images were compiled and batch-processed using the image browser with collective luminescent scales. Photon flux was measured as luminescent radiance (p/sec/cm^2^/sr). During luminescent threshold selection for image display, luminescent signals were regarded as background when minimum threshold levels resulted in displayed radiance above non-tissue-containing or known uninfected regions. To determine the pattern of virus spread, the image sequences were acquired every day following administration of SARS-CoV-2 (i.n). Image sequences were assembled and converted to videos using Image J.

#### Analyses of signature inflammatory cytokines mRNA

Brain and lung samples were collected from mice at the time of necropsy. Approximately, 20 mg of tissue was suspended in 500 µL of RLT lysis buffer, and RNA was extracted using RNeasy plus Mini kit (Qiagen Cat # 74136), reverse transcribed with iScript advanced cDNA kit (Bio-Rad Cat #1725036). To determine levels of signature inflammatory cytokines, multiplex qPCR was conducted using iQ Multiplex Powermix (Bio Rad Cat # 1725848) and PrimePCR Probe Assay mouse primers FAM-GAPDH, HEX-IL6, TEX615-CCL2, Cy5-CXCL10, and Cy5.5-IFNgamma. The reaction plate was analyzed using CFX96 touch real time PCR detection system. Scan mode was set to all channels. The PCR conditions were 95 °C 2 min, 40 cycles of 95 °C for 10 s and 60 °C for 45 s, followed by a melting curve analysis to ensure that each primer pair resulted in amplification of a single PCR product. mRNA levels of il6, ccl2, cxcl10 and ifng in the cDNA samples of infected mice were normalized to gapdh with the formula ΔC_t_(target gene)=C_t_(target gene)-C_t_(gapdh). The fold increase was determined using 2^-ΔΔCt^ method comparing treated mice to uninfected controls.

#### Protein expression and purification

FreeStyle 293F cells (Thermo Fisher) were grown in FreeStyle 293F medium (Thermo Fisher) to a density of 1x10^6^ cells/mL at 37°C with 8% CO_2_ with regular agitation (135 rpm). Cells were transfected with a plasmid coding for recombinant stabilized SARS-CoV-2 ectodomain (residue 1-1208) (S-6P or HexaPro Spike; obtained from Dr. Jason S. McLellan) (Hsieh et al., 2020), his-tagged SARS-CoV-2 RBD (residue 319-541) or SARS-CoV RBD-SD1(residue 306-577) using polyethylenimine (PEI). One-week post-transfection, supernatants were clarified and filtered using a 0.22 µm filter (Thermo Fisher Scientific). The crude S-6P was purified on strep-tactin resin (IBA) followed by size-exclusion chromatography on Superose 6 10/300 column (GE Healthcare) in 10 mM Tris pH 8.0 and 200 mM NaCl (SEC buffer). RBD was purified on a Ni-NTA column (Invitrogen) followed by size-exclusion chromatography on a Hiload 16/600 Superdex 200pg column using the same SEC buffer. For CryoEM protein sample preparation, the expression plasmid encoding S-6P was transfected into 293F GnT1- cells using PEI. The protein was harvested and purified with the same protocol as above. The C-terminal twin-Strep-Tag and 8xHis tag on S-6P was removed by HRV3C (Sigma) digestion as described in (Wrapp et al., 2020b) and the cleaved protein was further purified on a Ni-NTA column followed by gel filtration on a Superose 6 10/300 in SEC buffer. Purified proteins were snap-frozen in liquid nitrogen and stored in aliquots at -80°C until further use. Protein purity was confirmed by SDS-PAGE.

The expression plasmids encoding the heavy and light chains of CV3-13 IgG were transiently transfected into Expi293F cells (Thermo Fisher) using ExpiFectamine 293 transfection reagent as per the manufacturer’s protocol (Thermo Fisher). After 6-days post-transfection, the antibody was purified on Protein A resin from cell supernatant (Thermo Fisher) before the overnight papain digestion at 37°C using immobilized papain agarose (Thermo Fisher). The resulting Fab was separated from Fc and undigested IgG by passage over protein A resin. Fab was further purified by gel filtration using a Superose 6 10/300 column before use in SPR binding or cryo-EM sample preparation.

FreeStyle 293F cells (Thermo Fisher) were grown in FreeStyle 293F medium (Thermo Fisher Scientific) to a density of 1x10^6^ cells/mL at 37°C with 8% CO_2_ with regular agitation (135 rpm). The expression plasmids encoding the heavy and light chains of CV3-1, CV3-25, CV3-13 WT, CV3-13 LALA and CV3-13 GASDALIE IgG were transfected into Freestyle 293F cells (Thermo Fisher Scientific) using ExpiFectamine 293 transfection reagent as per the manufacturer’s protocol (Thermo Fisher Scientific). 1 week later, the cells were pelleted and discarded. The supernatants were filtered (Thermo Fisher Scientific) (0.22-μm-pore-size filter) as directed by the manufacturer (Thermo Fisher Scientific). The supernatant containing the antibody was then passed over Protein A beads (Cytiva). Following this, the antibodies were washed, eluted and dialyzed against PBS before their concentration being measured. Antibodies were then aliquoted and stored at -80°C until further use.

#### SARS-CoV-2 Spike ELISA (enzyme-linked immunosorbent assay)

The SARS-CoV-2 Spike ELISA assay used was recently described (Beaudoin-Bussieres et al., 2020; Prevost et al., 2020). Briefly, recombinant SARS-CoV-2 S-6P (2.5 μg/ml), or bovine serum albumin (BSA) (2.5 μg/ml) as a negative control, were prepared in PBS and were adsorbed to plates (MaxiSorp; Nunc) overnight at 4 °C. Coated wells were subsequently blocked with blocking buffer (Tris-buffered saline [TBS] containing 0.1% Tween20 and 2% BSA) for 1 hour at room temperature. Wells were then washed four times with washing buffer (TBS containing 0.1% Tween20). CV3-1, CV3-25, CV3-13 WT, CV3-13 LALA, CV3-13 GASDALIE and CR3022 mAbs (50 ng/ml) were prepared in a diluted solution of blocking buffer (0.1 % BSA) and incubated in the coated wells for 90 minutes at room temperature. Plates were washed four times with washing buffer followed by incubation with HRP-conjugated anti-Human IgG secondary Abs (Invitrogen) (at a concentration of 0.267 µg/mL in a diluted solution of blocking buffer [0.4% BSA]) for 1 hour at room temperature, followed by four washes. HRP enzyme activity was determined after the addition of 40 µL of a 1:1 mix of Western Lightning oxidizing and luminol reagents (Perkin Elmer Life Sciences). Light emission was measured with a LB941 TriStar luminometer (Berthold Technologies). Signal obtained with BSA was subtracted for each antibody and was then normalized to the signal obtained with CR3022 mAb present in each plate.

#### Flow cytometry analysis of cell-surface Spike staining

10 µg of the different expressors of the original SARS-CoV-2 Spike (Hoffmann et al., 2020) or the different mutants of the SARS-CoV-2 Spike (B.1.1.7, D614G, Δ69-70, Δ144, N501Y, A570D, P681H, T716I, S982A and D1118H) (Li et al., 2021b; Prevost et al., 2021; Tauzin et al., 2021) were co-transfected with 2.5 µg of a green fluorescent protein (GFP) expressor (pIRES2-eGFP) into 2 × 10^6^ 293T cells using the standard calcium phosphate method. Before staining with primary antibodies, cells were washed 2 times. At 48 hours post transfection, 293T cells were stained with CR3022, CV3-1, CV3- 13 WT, CV3-13 LALA, CV3-13 GASDALIE and CV3-25 antibodies (5μg/mL) for 45 minutes at 37°C before being washed 2 times in PBS. Alexa Fluor-647-conjugated goat anti-human IgG (H+L) Abs (Invitrogen) (2 µg/mL) and AquaVivid (Thermo Fischer Scientific) (Dilution of 1/1000) were used to stain the cells for 20 minutes at room temperature. The percentage of transfected cells (GFP+ cells) was determined by gating the living cell population based on the basis of viability dye staining (AquaVivid, Thermo Fischer Scientific). Samples were acquired on a LSRII cytometer (BD Biosciences) and data analysis was performed using FlowJo v10.5.3 (Tree Star).

For cell surface staining of transduced CEM.NKr-Spike cells, CEM.NKr-Spike cells were stained for 45 minutes at room temperature with CV3-1, CV3-13 WT, CV3-13 LALA and CV3-13 GASDALIE (0.0025 µg/mL, 0.01 µg/mL, 0.05 µg/mL, 0.25 µg/mL, µg/mL and 5 µg/mL) in PBS. Cells were then washed twice with PBS and stained with µg/mL of anti-human AlexaFluor 647 (AF-647) secondary antibody and 1:1000 dilution of viability dye AquaVivid (Thermo Fisher) for 20 minutes in PBS at room temperature. Cells were then washed twice with PBS and fixed in a 2% PBS-formaldehyde solution. All samples were acquired on an LSRII cytometer (BD Biosciences) and data analysis was performed using FlowJo v10.5.3 (Tree Star).

#### Surface plasmon resonance (SPR)

All surface plasma resonance assays were performed on a Biacore 3000 (GE Healthcare) with a running buffer of 10 mM HEPES pH 7.5 and 150 mM NaCl, supplemented with 0.05% Tween 20 at 25°C. Initial epitope mapping was performed by the binding of SARS-CoV RBD (residue 306-577) and other SARS-CoV-2 antigens (S1 and S2 obtained from BEI Resources) to immobilized CV3-13 IgG (∼5800 RU) on a Protein A sensor chip (Cytiva). For the kinetic measurement of CV3-13 Fab binding to SARS-CoV- 2 spike, ∼800 RU of his-tagged SARS-CoV-2 S-6P was immobilized on a Ni-pretreated NTA chip (Cytiva). 2-fold serial dilutions of purified CV3-13 Fab were then injected with concentrations ranging from 3.125 to 200 nM. Sensorgrams were corrected by subtraction of the corresponding blank channel as well as for the buffer background and kinetic constants determined using a 1:1 Langmuir model with the BIAevaluation software (GE Healthcare). Goodness of fit of the curve was evaluated by the Chi^2^ of the fit with a value below 3 considered acceptable.

#### Pseudovirus neutralization assay

Target cells were infected with single-round luciferase-expressing lentiviral particles. Briefly, 293T cells were transfected by the calcium phosphate method with the pNL4.3 R-E- Luc plasmid (NIH AIDS Reagent Program) and a plasmid encoding for SARS-CoV-2 Spike at a ratio of 10:1. Two days post-transfection, cell supernatants were harvested and stored at –80°C until use. 293T-ACE2 (Prevost et al., 2020) target cells were seeded at a density of 1 × 10^4^ cells/well in 96-well luminometer-compatible tissue culture plates (Perkin Elmer) 24 h before infection. Recombinant viruses in a final volume of 100 μL were incubated with the indicated semi-log diluted antibody concentrations (0 µg/mL, 0.01 µg/mL, 0.0316 µg/mL, 0.1 µg/mL, 0.316 µg/mL, 1 µg/mL, 3.16 µg/mL and 10 µg/mL) for 1 h at 37°C and were then added to the target cells followed by incubation for 48 h at 37°C; cells were lysed by the addition of 30 μL of passive lysis buffer (Promega) followed by one freeze-thaw cycle. An LB941 TriStar luminometer (Berthold Technologies) was used to measure the luciferase activity of each well after the addition of 100 μL of luciferin buffer (15 mM MgSO_4_, 15 mM KH_2_PO_4_ [pH 7.8], 1 mM ATP, and 1 mM dithiothreitol) and 50 μL of 1 mM d-luciferin potassium salt.

#### Microneutralization assay

One day prior to infection, 2x10^4^ Vero E6 cells were seeded per well of a 96 well flat bottom plate and incubated overnight to permit Vero E6 cell adherence. Antibody concentrations (0 µg/mL, 0.01 µg/mL, 0.0316 µg/mL, 0.1 µg/mL, 0.316 µg/mL, 1 µg/mL, 3.16 µg/mL and 10 µg/mL) were performed in a separate 96 well culture plate using DMEM supplemented with penicillin (100 U/mL), streptomycin (100 μg/mL), HEPES, 0.12% sodium bicarbonate, 2% FBS and 0.24% BSA. 10^4^ TCID_50_/mL of authentic SARS-CoV-2 D614G virus (derived from strain LSPQ/231457/2020; (Prevost et al., 2021)) was prepared in DMEM + 2% FBS and combined with an equivalent volume of diluted antibodies for one hour. After this incubation, all media was removed from the 96 well plate seeded with Vero E6 cells and virus:antibody mixture was added to each respective well at a volume corresponding to 600 TCID_50_ per well and incubated for one hour further at 37°C. Both virus only and media only (MEM + 2% FBS) conditions were included in this assay. All virus:antibody supernatant was removed from wells without disrupting the Vero E6 monolayer. Each diluted antibody (for a volume of 100 μL) was added to its respective Vero E6-seeded well in addition to an equivalent volume of MEM + 2% FBS and was then incubated for 48 hours. Media was then discarded and replaced with 10% formaldehyde for 24 hours to cross-link Vero E6 monolayer. After, formaldehyde was removed from wells and subsequently washed with PBS. Cell monolayers were permeabilized for 15 minutes at room temperature with PBS + 0.1% Triton X-100, washed with PBS and then incubated for one hour at room temperature with PBS + 3% non-fat milk. A mouse anti-SARS-CoV-2 nucleocapsid protein monoclonal antibody (Clone 1C7, Bioss Antibodies) solution was prepared at 1 μg/mL in PBS + 1% non-fat milk and added to all wells for one hour at room temperature. Following extensive washing (3×) with PBS, an anti-mouse IgG HRP secondary antibody solution was formulated in PBS + 1% non-fat milk. One hour post-room temperature incubation, wells were washed with 3×PBS, substrate (ECL) was added and an LB941 TriStar luminometer (Berthold Technologies) was used to measure the signal each well.

#### Cell-surface staining of SARS-CoV-2-infected cells

8M Vero-E6 cells were plated in T-175 flask 24 hours before infection. Authentic SARS-CoV-2 virus (MOI = 0.01) was used for infection. After 48 hours, infected Vero E6 cells were detached with PBS-EDTA and were incubated with 5 μg/mL of indicated antibodies for 30 minutes at 37℃, followed by staining with anti-human –AF647 secondary antibody and 1:1000 dilution of viability dye AquaVivid (Thermo Fisher Scientific) for 20 minutes at room temperature. Cells were then treated with 4% PFA for 24 hours at 4℃. Then the cells were stained intracellularly for SARS-CoV-2 nucleocapsid (N) antigen, using the Cytofix/Cytoperm fixation/permeabilization kit (BD Biosciences) and an anti-N mAb (clone mBG17; Kerafast) conjugated with the Alexa Fluor 488 dye according to the manufacturer’s instructions (Invitrogen). The percentage of infected cells (N+ cells) was determined by gating the living cell population based on the basis of viability dye staining (Aqua Vivid, Invitrogen). Samples were acquired on a LSRII cytometer (BD Biosciences, Mississauga, ON, Canada) and data analysis was performed using FlowJo v10.5.3 (Tree Star, Ashland, OR, USA).

#### Antibody dependent cellular cytotoxicity (ADCC) assay

This assay was previously described (Anand et al., 2021). Briefly, for evaluation of anti-SARS-CoV-2 ADCC activity, parental CEM.NKr CCR5+ cells were mixed at a 1:1 ratio with CEM.NKr-Spike cells. These cells were stained for viability (AquaVivid; Thermo Fisher Scientific) and a cellular dye (cell proliferation dye eFluor670; Thermo Fisher Scientific) and subsequently used as target cells. Overnight rested PBMCs were stained with another cellular marker (cell proliferation dye eFluor450; Thermo Fisher Scientific) and used as effector cells. Stained effector and target cells were mixed at a 10:1 ratio in 96-well V-bottom plates. Titrated concentrations of CV3-1, CV3-13 WT, CV3-13 LALA and CV3-13 GASDALIE monoclonal antibodies (0.0025 µg/mL, 0.01 µg/mL, 0.05 µg/mL, 0.25 µg/mL, 1 µg/mL and 5 µg/mL) were added to the appropriate wells. The plates were subsequently centrifuged for 1 min at 300 x g, and incubated at 37°C, 5% CO_2_ for 5 hours before being fixed in a 2% PBS-formaldehyde solution. Since CEM.NKr-Spike cells express GFP, ADCC activity was calculated using the formula: [(% of GFP+ cells in Targets plus Effectors) - (% of GFP+ cells in Targets plus Effectors plus antibody)] / (% of GFP+ cells in Targets) x 100 by gating on transduced live target cells. All samples were acquired on an LSRII cytometer (BD Biosciences) and data analysis performed using FlowJo v10.5.3 (Tree Star).

#### Antibody dependent cellular phagocytosis (ADCP) assay

The ADCP assay was performed using CEM.NKr-Spike cells as target cells that were fluorescently labelled with a cellular dye (cell proliferation dye eFluor450). THP-1 cells were used as effector cells and were stained with another cellular dye (cell proliferation dye eFluor670). Stained target cells were incubated with 100 µL of the titrated concentrations of CV3-1, CV3-13 WT, CV3-13 LALA and CV3-13 GASDALIE antibodies (0,625 µg/mL, 1,25 µg/mL, 2,5 µg/mL, 5 µg/mL and 10 µg/mL) for 1h at 37°C, followed by two washes with media. Stained target and effector cells were mixed at a 5:1 ratio in 96-well U-bottom plates. After a 5 hours incubation at 37 °C and 5% CO_2_, cells were fixed with a 2% PBS-formaldehyde solution. Antibody-mediated phagocytosis was determined by flow cytometry, gating on THP-1 cells that were triple-positive for eFluor450 and eFluor670 cellular dyes and GFP. All samples were acquired on an LSRII cytometer (BD Biosciences) and data analysis performed using FlowJo v10.5.3 (Tree Star).

#### Cryo-EM sample preparation and data collection

SARS-CoV-2 HexaPro spike (GnT1- produced) was incubated with 20-fold excess of CV3-13 Fab overnight at 4°C before purification on a Superose 6 300/10 GL column (GE Healthcare). The complex peak was harvested, concentrated to 0.50 mg/mL in the SEC buffer and immediately used for Cryo-EM grid preparation. 3μL of protein was deposited on a holey copper grids (QUANTIFOIL R 1.2/1.3, 200 mesh, EMS) which had been glow-discharged for 30s at 15 Ma using PELCO easiGlow (TedPella Inc). Grids were vitrified in liquid ethane using Vitrobot Mark IV (Thermo Fisher) with a blot time of 2-4 s and variable blot force at 4°C and 100% humidity.

The frozen grids were screened on a FEI Talos Arctica microscope at 200 kV equipped with a FEI Falcon3EC detector using the EPU software (Thermo Fisher). Cryo-EM data from a good grid were acquired on a FEI Glacios electron microscope operating at 200 kV, equipped with a Gatan K3 direct electron detector. Micrographs were collected at a magnification of 45,000 corresponding to a calibrated pixel size of 0.8893 Å, with a total exposure dose of 42 e-/ Å^2^.

#### Cryo-EM data processing, model building and analysis

Motion correction, CTF estimation, particle picking, curation and extraction, 2D classification, *ab initio* model reconstruction, volume refinements and local resolution estimation were carried out in cryoSPARC (Punjani et al., 2017; Rubinstein and Brubaker, 2015). An initial SARS-CoV-2 spike model (PDB: 6XKL) with single-RBD up and a three-RBD-down (closed) model (PDB: 6VXX) were used as modeling templates. The NTDs were initially modelled from PDB entry 7C2L. The initial models for CV3-13 Fab were generated in the SAbPred server (Dunbar et al., 2016) using the primary sequence.

Automated and manual model refinements were iteratively carried out in ccpEM (Burnley et al., 2017), Phenix (Liebschner et al., 2019) (real-space refinement) and Coot (Emsley and Cowtan, 2004). Geometry validation and structure quality evaluation were performed by EM-Ringer (Barad et al., 2015) and Molprobity (Chen et al., 2010). Model-to-map fitting cross correlation and figures generation were carried out in USCF Chimera, Chimera X (Pettersen et al., 2021) and PyMOL (The PyMOL Molecular Graphics System, Version 1.2r3pre, Schrödinger, LLC). The complete cryo-EM data processing workflow is shown in Figure S4 and statistics of data collection, reconstruction and refinement is described in Table S1. The epitope interface analysis was performed in PISA (Krissinel and Henrick, 2007).

#### Quantification and Statistical Analysis

Statistical significance was derived by applying parametric unpaired t-test or non-parametric Mann-Whitney U test (two-tailed) available in GraphPad Prism software (La Jolla, CA, USA) depending on the normality distribution of the data. *p* values lower than 0.05 were considered statistically significant. P values were indicated as ∗, p < 0.05; ∗∗, p < 0.01; ∗∗∗, p < 0.001; ∗∗∗∗, p < 0.0001.

## Supplementary Figure Legends

**Figure S1.**
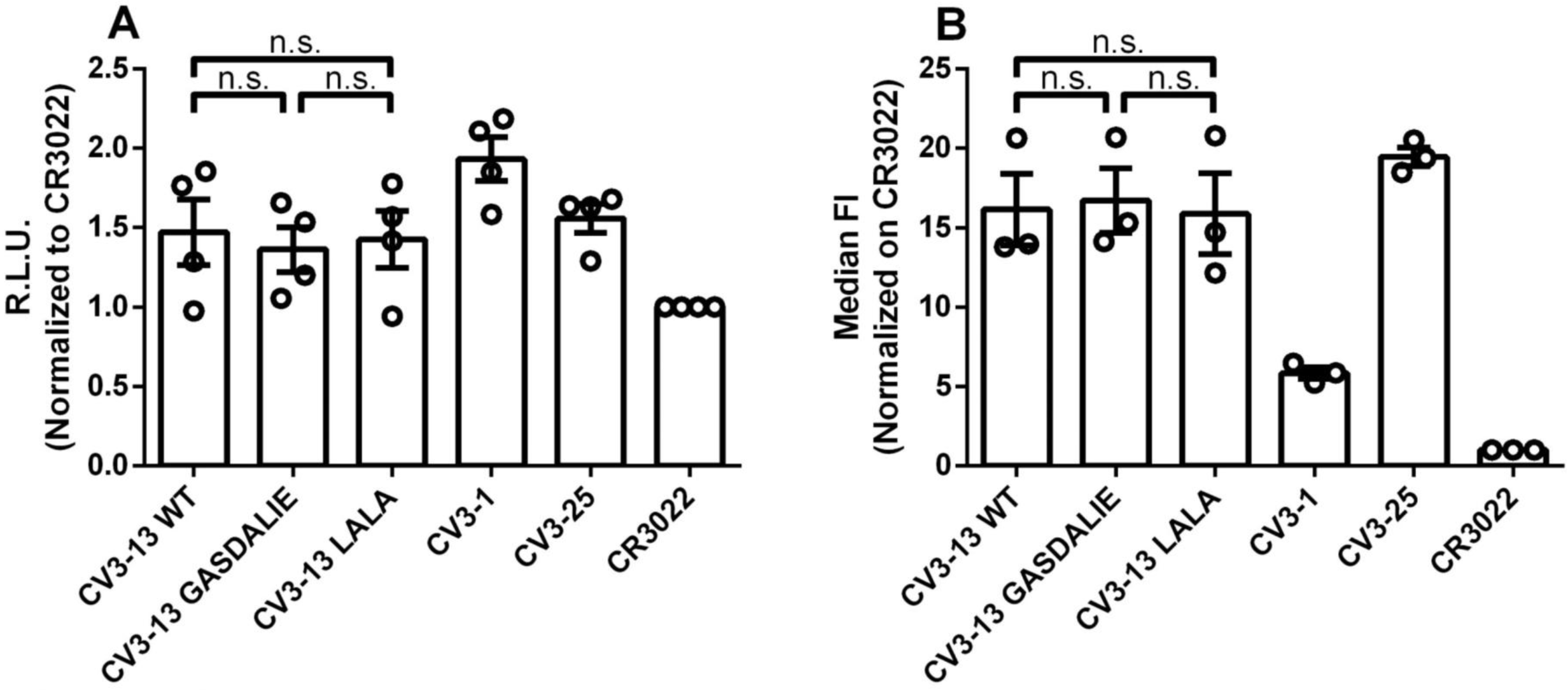
LALA and GASDALIE Fc mutations does not affect CV3-13 binding to SARS-CoV-2 Spike. (A) Indirect ELISA on SARS-CoV-2 Spike 6P using CV3-13 WT, CV3-13 LALA, CV3- 13 GASDALIE, CV3-1, CV3-25 and CR3022 (50 ng/mL). CR3022 was used as a positive control in each ELISA plate and for each experiment the data was further normalized on the signal obtained with this antibody. Experiment was done 4 times. (B) Staining with CV3-13 WT, CV3-13 LALA, CV3-13 GASDALIE, CV3-1, CV3-25 and CR3022 (5 µg/mL) of 293T cells transfected with the SARS-CoV-2 Spike. CR3022 was used as a positive control in each experiment and for each experiment the data was further normalized on the signal obtained with this antibody. Experiment was done 3 times. Statistical significance was evaluated using a non-parametric Mann-Whitney U test (n.s., not significant). The data shown is the mean ± SEM.

**Figure S2.**
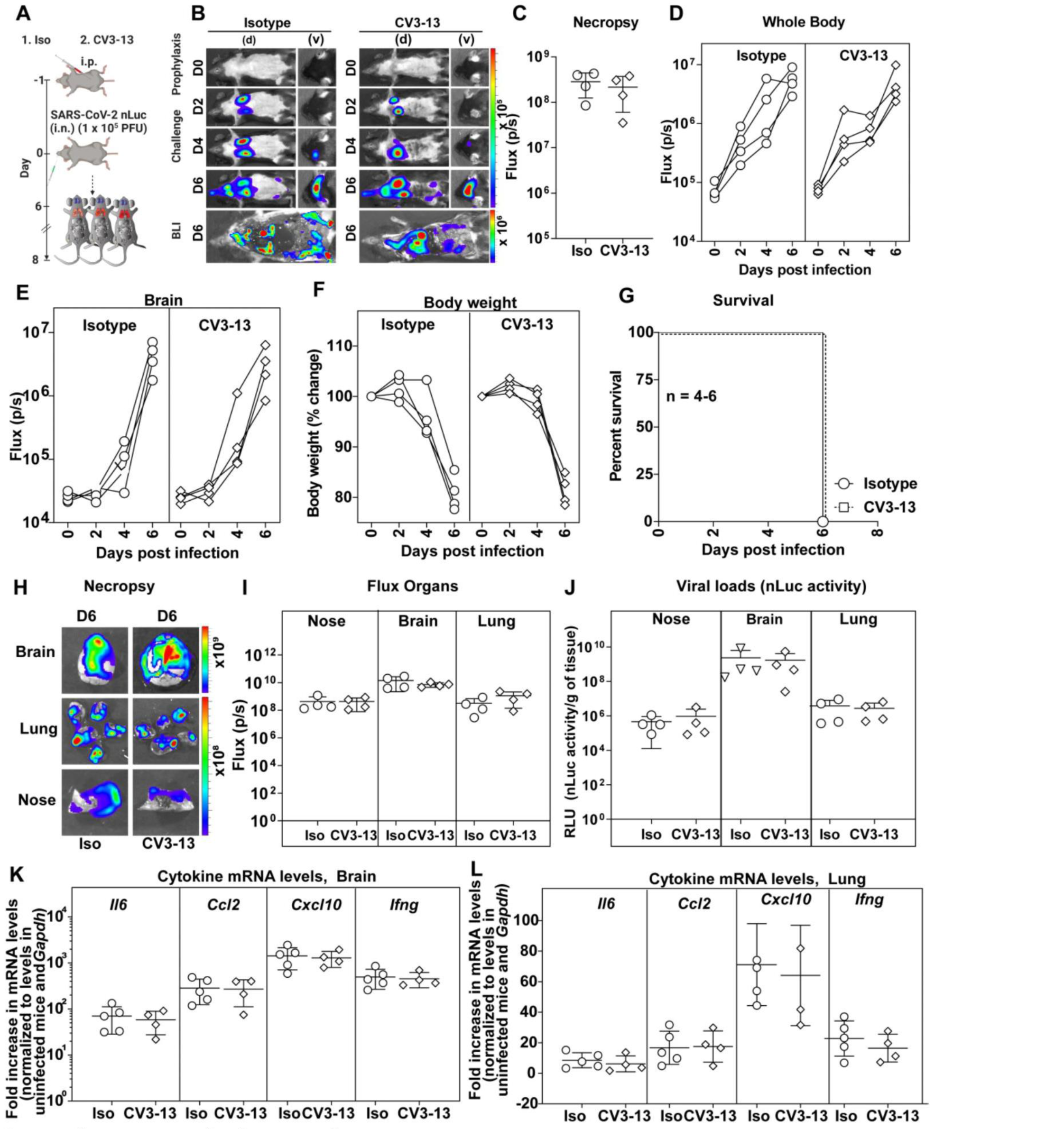
Prophylactic treatment with non-neutralizing CV3-13 antibody does not protect K18-hACE2 mice from lethal SARS-CoV-2 infection. (A) Experimental design for testing *in vivo* efficacy of non-neutralizing CV3-13 antibody administered 1 day prior to challenging K18-hACE2 mice (i.n.) with SARS-CoV-2-nLuc followed by non-invasive BLI every 2 days. Human IgG1-treated (12.5 mg IgG/kg) mice were use as the isotype control (Iso). (B) Representative images from BLI of SARS-CoV- 2-nLuc-infected mice in ventral (v) and dorsal (d) positions at the indicated dpi and after necropsy for experiment as in A. (C) *Ex-vivo* quantification of nLuc signal as flux (photons/sec) after necropsy. (D, E) Temporal quantification of nLuc signal as flux (photons/sec) computed non-invasively in indicated areas of each animal. (F) Temporal changes in mouse body weight with initial body weight set to 100 %. (G) Kaplan-Meier survival curves of mice statistically compared by log-rank (Mantel-Cox) test for experiment as in A. (H, I) *Ex-vivo* imaging of organs and quantification of nLuc signal as flux (photons/sec) at the indicated dpi after necropsy. (J) Viral loads (nLuc activity/g) from indicated organs using Vero E6 cells as targets. (K, L) Cytokine mRNA levels in lung and brain tissues after necropsy normalized to *Gapdh* in the same sample and that in uninfected mice. Viral loads (J) and inflammatory cytokine profile (K, L) were determined after necropsy for mice that succumbed to infection. Scale bars in (B) and (H) denote radiance (photons/sec/cm^2^/steradian). Each curve in (D)-(F) and each data point in (C) and (I)-(L) represents an individual mouse. The data in (C) and (I)-(L) were analyzed by non-parametric Mann Whitney test. Non-significant values are not shown, Mean values ± SD are depicted.

**Figure S3.**
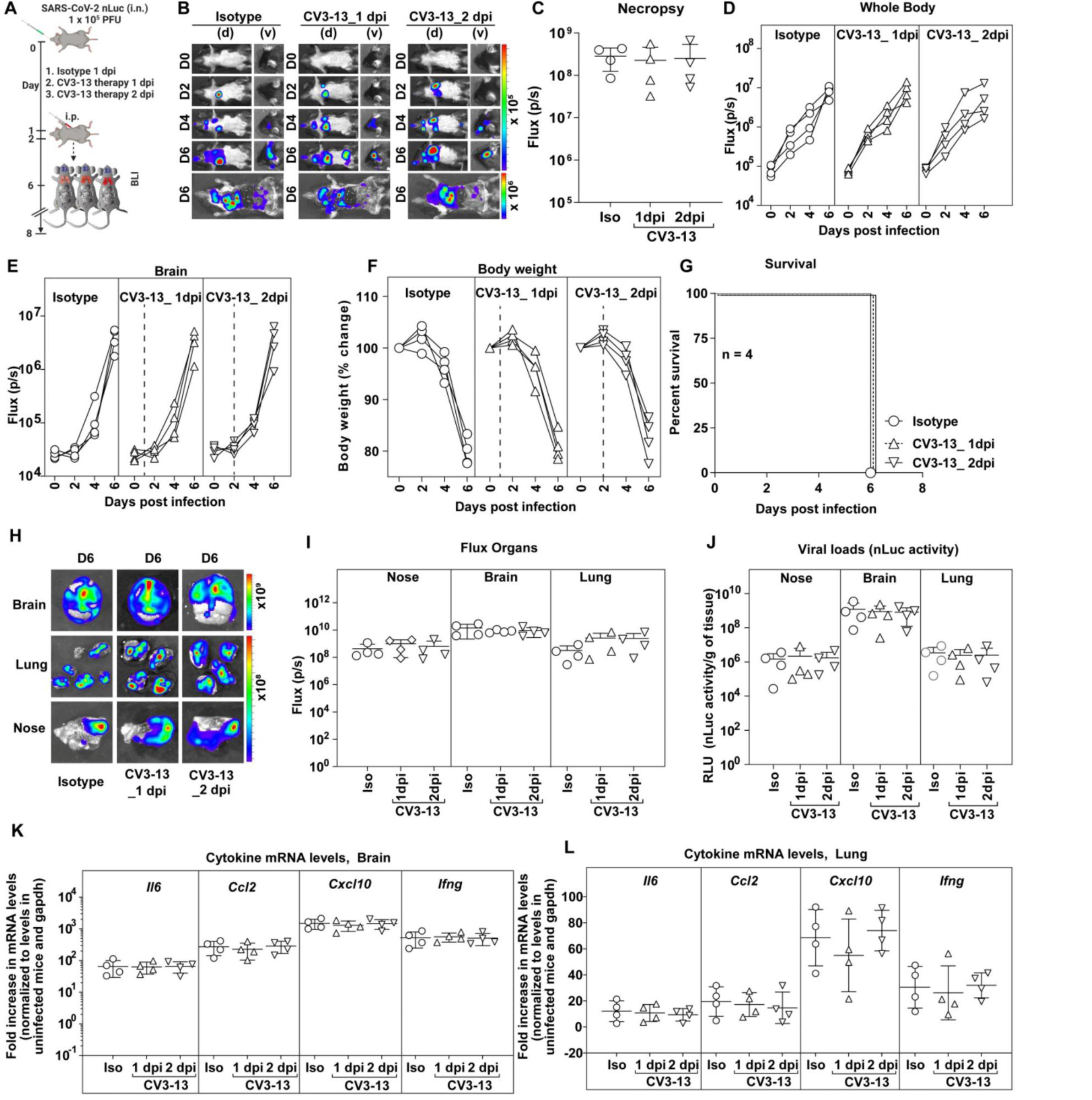
Therapeutic treatment with non-neutralizing CV3-13 antibody does not protect K18-hACE2 mice from lethal SARS-CoV-2 infection. (A) Experimental design for testing *in vivo* efficacy of CV3-13 administered at 1 and 2 dpi after challenging K18-hACE2 mice (i.n.) with SARS-CoV-2-nLuc followed by non- invasive BLI every 2 days. Human IgG1-treated (12.5 mg IgG/kg) mice were use as the isotype control (Iso). (B) Representative images from BLI of SARS-CoV-2-nLuc- infected mice in ventral (v) and dorsal (d) positions at the indicated dpi and after necropsy at indicated days for experiment as in A. (C) *Ex-vivo* quantification of nLuc signal as flux (photons/sec) after necropsy. (D, E) Temporal quantification of nLuc signal as flux (photons/sec) computed non-invasively in indicated areas of each animal. (F) Temporal changes in mouse body weight with initial body weight set to 100 %. (G) Kaplan-Meier survival curves of mice statistically compared by log-rank (Mantel-Cox) test for experiment as in A. (H, I) *Ex-vivo* imaging of organs and quantification of nLuc signal as flux (photons/sec) at the indicated dpi after necropsy. (J) Viral loads (nLuc activity/g) from indicated organs using Vero E6 cells as targets. (K, L) Cytokine mRNA levels in lung and brain tissues after necropsy normalized to *Gapdh* in the same sample and that in uninfected mice. Viral loads (J) and inflammatory cytokine profile (K, L) were determined after necropsy for mice that succumbed to infection. Scale bars in (B) and (H) denote radiance (photons/sec/cm^2^/steradian). Each curve in (D)-(F) and each data point in (C) and (I)-(L) represents an individual mouse. The data in (C) and (I)-(L) were analyzed by Mann Whitney U test. Mean values ± SD are depicted.

**Figure S4.**
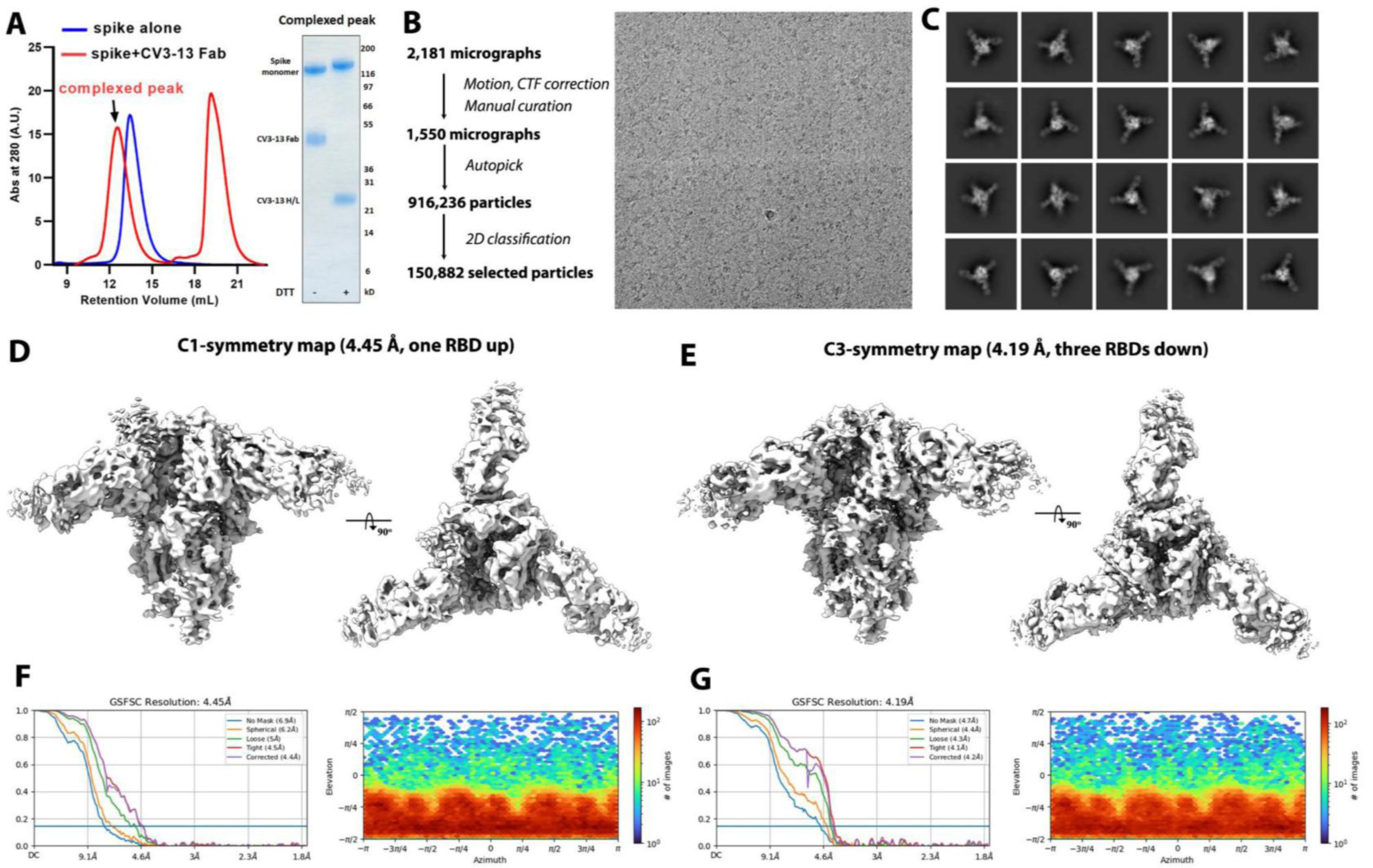
Cryo-EM data collection and processing of SARS-CoV-2 HexaPro Spike-CV3-13 Fab complex. (A) Cryo-EM sample preparation. Size-exclusion chromatogram (SEC) of the purified, non-tagged SARS-CoV-2 Hexa-Pro spike (blue) and its overnight mixture with CV3-13 Fab (molar-ratio: 1:20) (red) on a superpose 6 300/10 GL column. The SEC peak-shift and SDS-PAGE validate the Fab-spike complex formation. (B) Cryo-EM data processing workflow in cryosparc and the representative raw electron micrograph. (C) Selected 2D averages for *ab initio* reconstruction. (D, E) Side and top views of the final cryo-EM density map imposed with C1 (D) or C3 symmetry (E). (F, G) The Fourier shell correlation curves indicate the overall resolution (FSC cutoff 0.143) using non-uniform refinement (left panel) and the direction distribution plot of all particles used in the final refinement (right panel) for the C1-symmetry map (F) and the C3-symmetry map (G).

**Figure S5.**
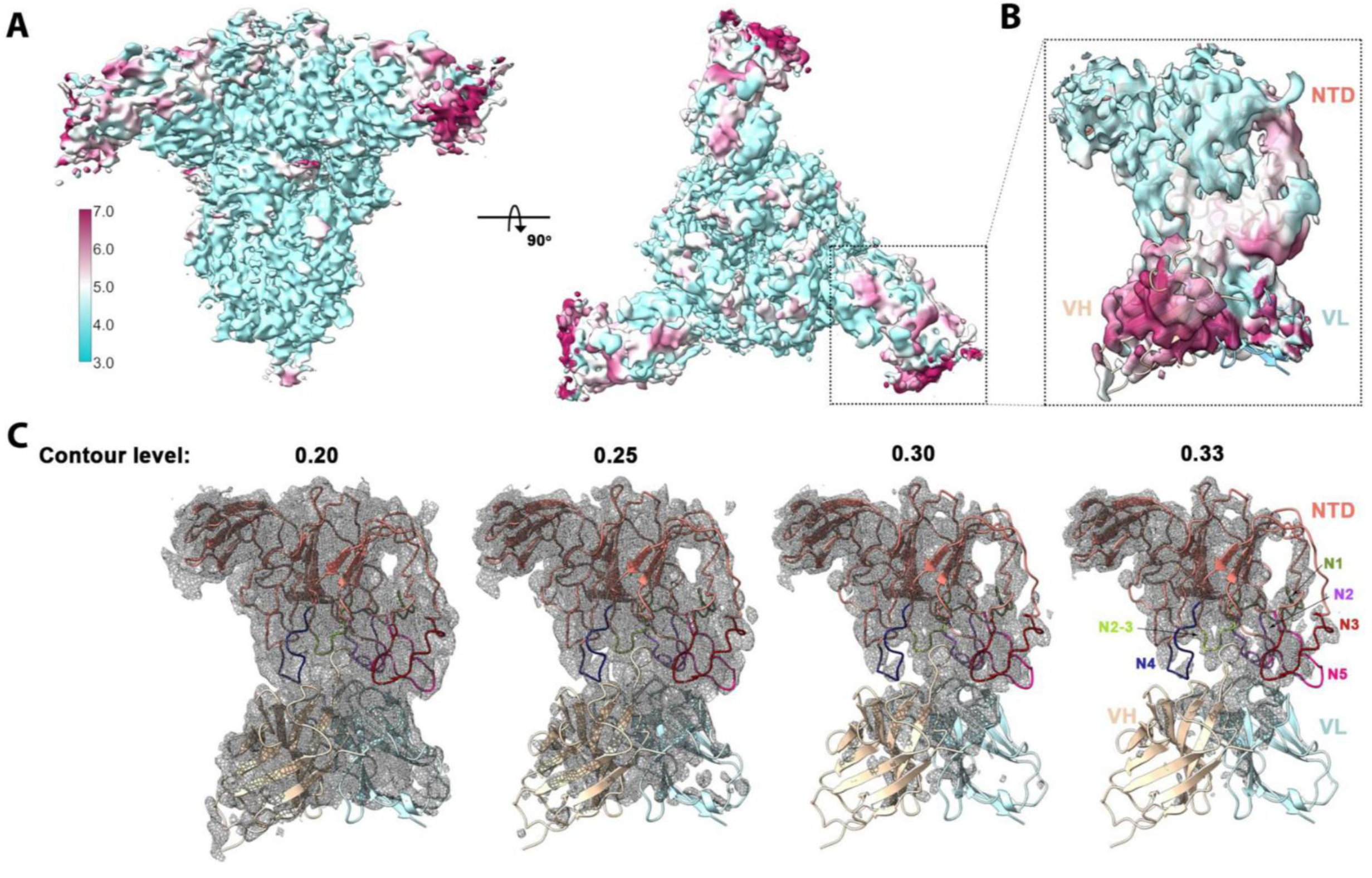
Cryo-EM structure validation. (A) Two views of C3-symmetry density map colored by local resolution as calculated in cryoSPARC using a FSC cutoff of 0.143. (B) Blow-up view of the density corresponding to the NTD-Fv portion colored by local resolution. (C) Density fit of NTD and CV3-13 Fv at different contour levels (Chimera X) in which the NTD framework is shown as salmon ribbons with NTD loops colored as indicated and the heavy and light chain of CV3-13 variable region colored in light yellow and cyan.

**Figure S6.**
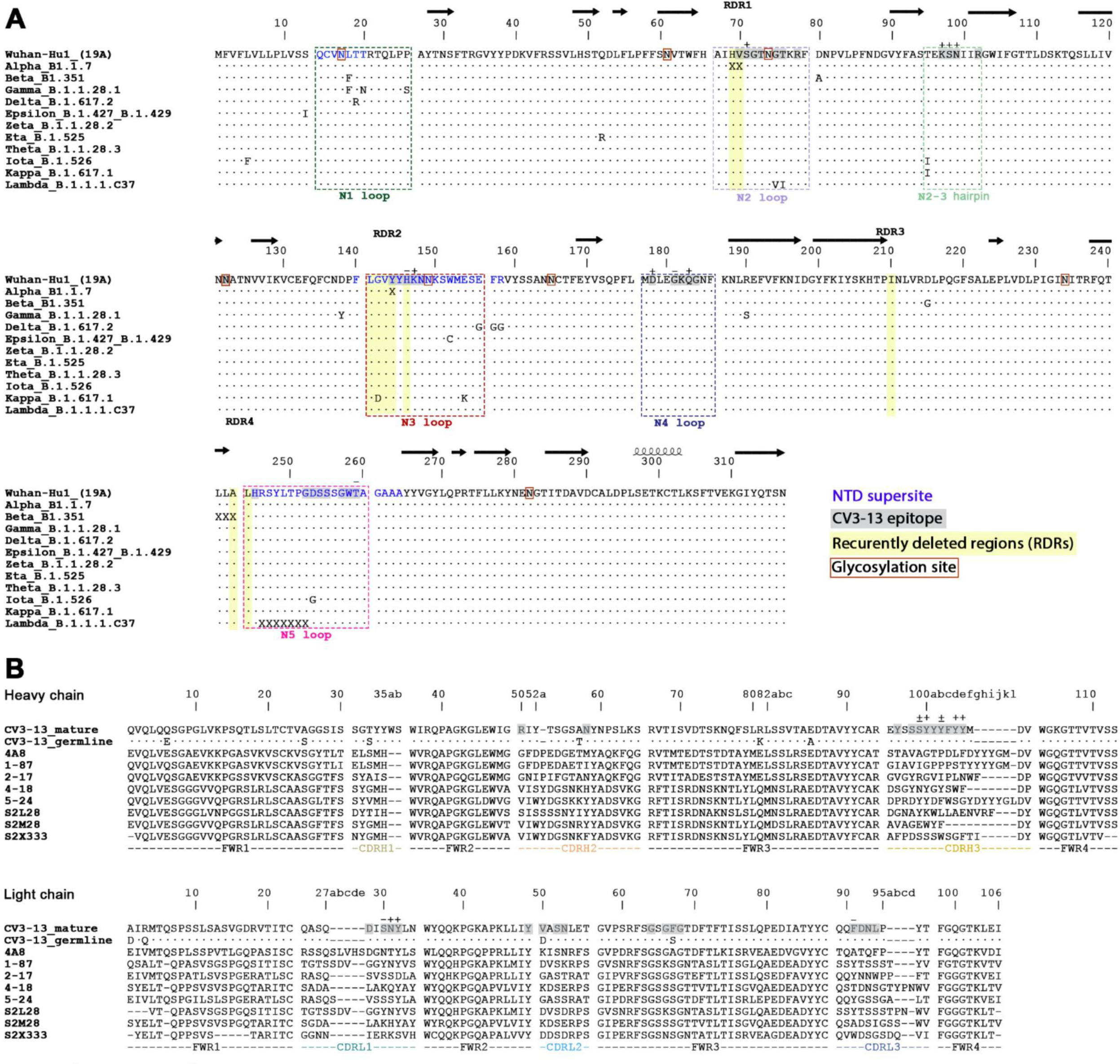
Sequence alignments. (A) Sequence alignment of the NTD (residue 1-317) from 11 SARS-CoV-2 variants. The sequence of ancestral Wuhan strain is listed on the top and the mutations from other variants are listed (“X” denotes a deletion), while the remaining identical residues to the Wuhan strain are shown as “·” for clarify. CV3-13 contacting residues are shaded in grey with the hydrogen-bonded residues marked above the sequence (“+” for the side chain and “-” for the main chain). Residues in the NTD supersite are colored in blue and the recurrently deleted regions (RDR1-4), as described by McCathy and colleagues (McCarthy et al., 2021), are shaded in yellow. The regions corresponding to N1-N5 loops and the N2-3 hairpin are highlighted in colored dashed rectangles. The documented glycosylation sites are boxed in red. The secondary structure assignment on top is derived from the 4A8-spike complex (PDB: 7C2L), the first reported NTD-directed mAb-Spike structure (Chi et al., 2020). (B) V_H_ and V_L_ sequence alignments of the affinity-matured CV3-13 and its germline IGHV4 and IGKV1 and the other NTD-directed mAbs. The NTD-contact residues are shaded in grey with those involved in hydrogen-bonds marked with (+) for side chain and (-) for mainchain above the sequence. The somatically mutated residues in the germline sequence are as shown with the unchanged residues marked as “·”.

**Supplemental table 1.** Cryo-EM data collection and refinement statistics.

| Protein | SARS-CoV-2 HexaPro spike/CV3-13 Fab |
| --- | --- |
| EMDB | EMD-24628 |
| PDB | 7RQ6 |
| **Data collection and Reconstruction** |  |
| Microscope | FEI Glacios |
| Voltage (kV) | 200 |
| Electron dose (e<sup>-</sup>/Å<sup>2</sup>) | 42 |
| Detector | Gatan K3 |
| Magnification | 45,000 |
| Pixel size (Å/pix) | 0.889 |
| Defocus range (µm) | 0.27-3.03 |
| Micrographs collected | 2,181 |
| Particles extracted/final | 916,236/98,752 |
| Symmetry imposed | C3 |
| Box size (pix) | 512 |
| Unmasked resolution at 0.5/0.143 FSC (Å) | 4.45 / 4.22 |
| Masked resolution at 0.5/0.143 FSC (Å) | 4.37 /4.19 |
| **Refinement and Validation** |  |
| Software | CryoSPARC/Phenix 1.18/ccpEM |
| Protein residues | 3951 |
| Chimera CC | 0.868 |
| EMRinger Score | 1.77 |
| Model vs data CC | 0.76 |
| R.m.s. Deviations |  |
| Bond Length (Å) | 0.004 |
| Bond Angles (°) | 0.726 |
| Molprobability score | 2.14 |
| Clash score | 13.20 |
| Favored rotamers (%) | 99.94 |
| Ramamchandran |  |
| Favored regions (%) | 91.34 |
| Allowed regions (%) | 8.58 |
| Disallowed regions (%) | 0.08 |

